# *De novo* cobalamin biosynthesis, transport and assimilation and cobalamin-mediated regulation of methionine biosynthesis in *Mycobacterium smegmatis*

**DOI:** 10.1101/2020.11.04.369181

**Authors:** Terry Kipkorir, Gabriel T. Mashabela, Timothy J. De Wet, Anastasia Koch, Lubbe Wiesner, Valerie Mizrahi, Digby F. Warner

**Affiliations:** SAMRC/NHLS/UCT Molecular Mycobacteriology Research Unit, DSI/NRF Centre of Excellence for Biomedical TB Research, Department of Pathology and Institute of Infectious Disease and Molecular Medicine, Faculty of Health Sciences, University of Cape Town, Observatory 7925, South Africa; Department of Integrative Biomedical Sciences, Faculty of Health Sciences, University of Cape Town, Observatory 7925, South Africa; Division of Clinical Pharmacology, Department of Medicine, University of Cape Town, Cape Town, South Africa; Wellcome Centre for Infectious Diseases Research in Africa, Faculty of Health Sciences, University of Cape Town, Observatory 7925, South Africa

**Keywords:** vitamin B_12_, riboswitch, *cobK*, tuberculosis

## Abstract

Cobalamin is an essential co-factor in all domains of life, yet its biosynthesis is restricted to some bacteria and archaea. *Mycobacterium smegmatis*, an environmental saprophyte frequently used as surrogate for the obligate human pathogen, *M. tuberculosis*, carries approximately 30 genes predicted to be involved in *de novo* cobalamin biosynthesis. *M. smegmatis* also encodes multiple cobalamin-dependent enzymes, including MetH, a methionine synthase which catalyses the final reaction in methionine biosynthesis. In addition to *metH*, *M. smegmatis* possesses a cobalamin-independent methionine synthase, *metE*, suggesting that enzyme selection – MetH or MetE – is regulated by cobalamin availability. Consistent with this notion, we previously described a cobalamin-sensing riboswitch controlling *metE* expression in *M. tuberculosis*. Here, we apply a targeted mass spectrometry-based approach to confirm *de novo* cobalamin biosynthesis in *M. smegmatis* during aerobic growth *in vitro*. We also demonstrate that *M. smegmatis* transports and assimilates exogenous cyanocobalamin (CNCbl; a.k.a. vitamin B_12_) and its precursor, dicyanocobinamide ((CN)_2_Cbi). Interestingly, the uptake of CNCbl and (CN)_2_Cbi appears restricted in *M. smegmatis* and dependent on the conditional essentiality of the cobalamin-dependent methionine synthase. Using gene and protein expression analyses combined with single-cell growth kinetics and live-cell time-lapse microscopy, we show that transcription and translation of *metE* are strongly attenuated by endogenous cobalamin. These results support the inference that *metH* essentiality in *M. smegmatis* results from riboswitch-mediated repression of MetE expression. Moreover, differences observed in cobalamin-dependent metabolism between *M. smegmatis* and *M. tuberculosis* provide some insight into the selective pressures which might have shaped mycobacterial metabolism for pathogenicity.

**IMPORTANCE:** Accumulating evidence suggests that alterations in cobalamin-dependent metabolism marked the evolution of *Mycobacterium tuberculosis* from an environmental ancestor to an obligate human pathogen. However, the roles of cobalamin in mycobacterial physiology and pathogenicity remain poorly understood. We used the non-pathogenic saprophyte, *M. smegmatis*, to investigate the production of cobalamin, transport and assimilation of cobalamin precursors, and the potential role of cobalamin in regulating methionine biosynthesis. We provide biochemical and genetic evidence confirming constitutive *de novo* cobalamin biosynthesis in *M. smegmatis* under standard laboratory conditions, in contrast with *M. tuberculosis*, which appears to lack *de novo* cobalamin biosynthetic capacity. We also demonstrate that the uptake of cyanocobalamin (vitamin B_12_) and its precursors is restricted in *M. smegmatis*, apparently depending on the need to service the co-factor requirements of the cobalamin-dependent methionine synthase. These observations support the utility of *M. smegmatis* as a model to elucidate key metabolic adaptations enabling mycobacterial pathogenicity.

## INTRODUCTION

Several mycobacterial species have been identified among the subset of prokaryotes which possess the genetic capacity for *de novo* cobalamin biosynthesis (1–5).

Included in this list of potential cobalamin producers is *Mycobacterium smegmatis*, the saprophytic mycobacterium commonly used as a surrogate for *Mycobacterium tuberculosis*, which causes tuberculosis (TB), a deadly respiratory disease claiming over 1 million lives globally every year (6–8). Cobalamin has one of the most complex structures of any of the biological cofactors, comprising a tetrapyrrole framework with a centrally chelated cobalt ion, a lower axial base (α ligand) which is typically dimethylbenzimidazole (DMB), and an upper axial ligand (R-group; β ligand) (Figure 1A). The nomenclature and catalytic activity of cobalamin depends on the β ligand. For example, in adenosylcobalamin (AdoCbl; a.k.a. coenzyme B_12_), the β ligand is a deoxyadenosyl group utilised by isomerases such as the methylmalonyl-CoA mutase and class II ribonucleotide reductases. In methylcobalamin (MeCbl), which is used by methyltransferases such as methionine synthase, the β ligand is a methyl group (Figure 1A) (9, 10).

**Figure 1.**
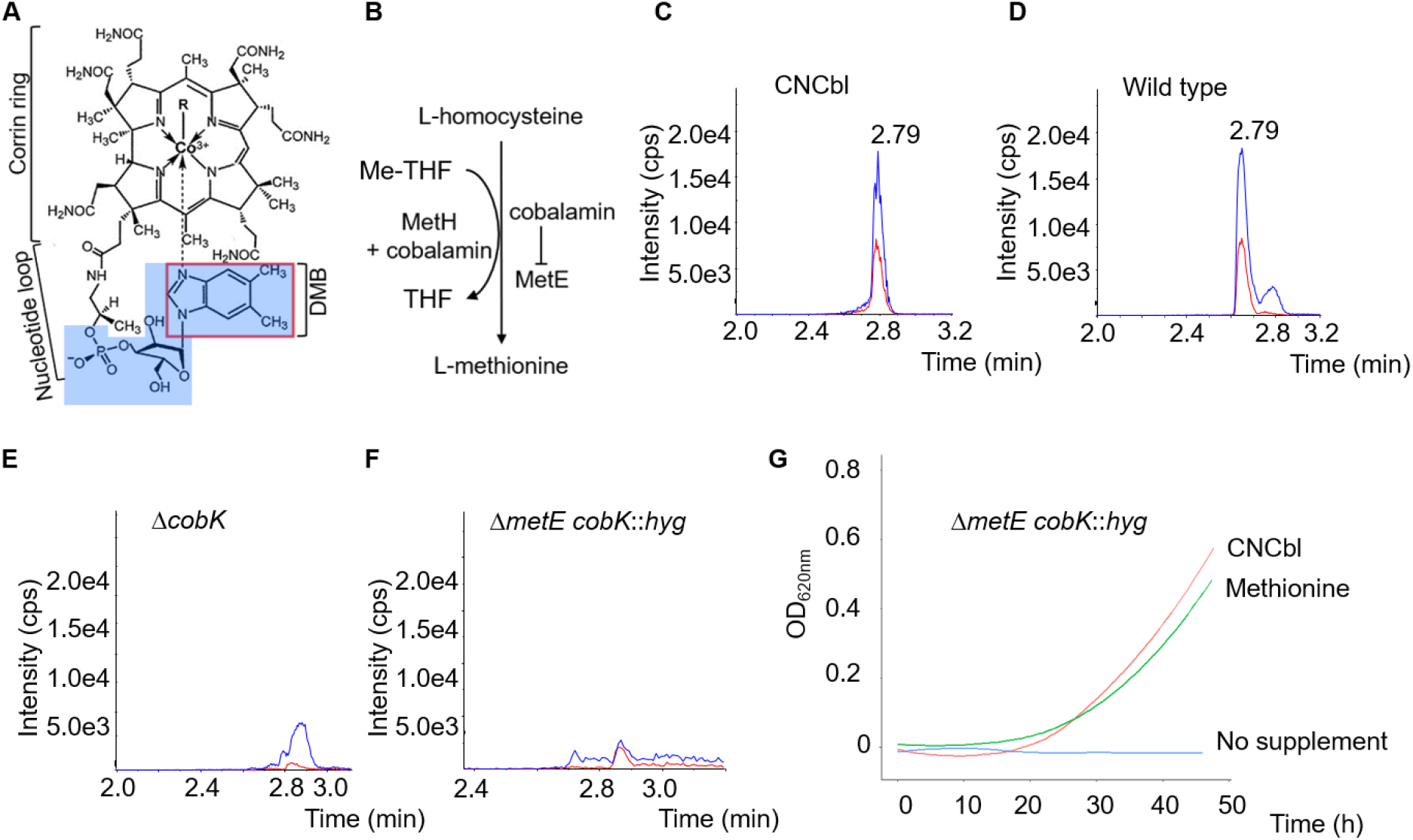
*De novo* cobalamin biosynthesis in *M. smegmatis*. **A**. Cobalamin structure. The cobalt ion is coordinated equatorially by four nitrogen atoms of a corrin ring and axially by variable lower (α) and upper (β) ligands (R-group). Examples of β ligands are CN in cyanocobalamin (CNCbl; a.k.a. vitamin B_12_), adenosyl in adenosylcobalamin (AdoCbl; a.k.a. coenzyme B_12_) and methyl in methylcobalamin (MeCbl). The α ligand in the physiologically relevant cobalamin is typically dimethylbenzimidazole (DMB; outlined in red box). **B.** The final step in the methionine biosynthesis pathway is a non-reversible transfer of a methyl group from methyltetrahydrofolate (Me-THF) to homocysteine to produce methionine and tetrahydrofolate (THF). This reaction is catalysed by either MetH using cobalamin as co-factor, or MetE. MetE expression is attenuated by cobalamin via a cobalamin sensing riboswitch. **C.** The LC-MS/MS method optimised to detect co-eluting peaks corresponding to α-ribazole 5’-phosphate (highlighted in blue shade in ***A***; blue trace) and DMB (red trace) transitions in a 20ng/ml CNCbl standard. **D-F.** Detection of *de novo* derivatised CNCbl. Cobalamin was detected in wild type (**D**) but not in Δ*cobK* (**E**) nor Δ*metE cobK*::*hyg* (**F**) mutants. Peak intensities are expressed as counts per second (cps). **G.** Growth curves of the Δ*metE cobK*::*hyg* strain in liquid 7H9-OADC medium in the presence of 10μM CNCbl or 1mM methionine. The mutant cannot grow without supplementation.

Cobalamin biosynthesis is a complex multi-step process requiring nearly thirty enzyme-catalysed biotransformations including eight SAM-dependent methylations, ring contraction, six amidations, decarboxylation, cobalt insertion, aminopropanol attachment and the assembly and attachment of the α ligand (11). Owing to the heavy energetic investment necessary to support the *de novo* pathway, this process is typically augmented in many organisms by the capacity for uptake and salvage (1). Interestingly, cobalamin-producing bacteria encode a much larger complement of genes involved in biosynthesis and salvage than the number of cobalamin-dependent metabolic pathways in those organisms. The preservation of *de novo* biosynthetic capacity therefore suggests the contribution of cobalamin to adaptation to specific lifestyles – an interpretation which is especially intriguing in the context of pathogenic, parasitic and symbiotic bacteria (4).

Like most mycobacteria, *M. smegmatis* encodes several cobalamin-dependent enzymes (Table 1) (12), some of which appear redundant given the existence of isoenzymes or alternative mechanisms for the same metabolic pathway (13). Among these are the methionine synthases, MetH (5-methyltetrahydrofolate-homocysteine methyltransferase, EC: 2.1.1.14) and MetE (5-methyltetrahydropteroyltriglutamate-homocysteine methyltransferase, EC: 2.1.1.13), which catalyse the non-reversible transfer of a methyl group from 5-methyltetrahydrofolate to homocysteine in the final step in the biosynthesis of methionine, an essential amino acid required for translation initiation, DNA methylation and cysteine biosynthesis (14, 15). MetH requires cobalamin for activity while MetE is a cobalamin-independent methionine synthase (Figure 1B) (9, 16, 17). We previously demonstrated that a cobalamin-sensing riboswitch located in the 5’ untranslated region (5’ UTR) of the *metE* gene in *M. tuberculosis* attenuated *metE* transcript levels in the presence of exogenous cyanocobalamin (CNCbl; a.k.a. vitamin B_12_) (18). We also showed that *M. tuberculosis* CDC1551, a well-characterised strain isolated during a TB outbreak in the US (19), contains a natural truncation of the *metH* gene which renders the strain sensitive to exogenous CNCbl. This phenotype, which was recapitulated in an engineered *M. tuberculosis* H37Rv mutant containing an analogous *metH* truncation, suggested that the observed growth inhibition was due to methionine depletion resulting from the effective elimination of all methionine synthase activity in the CNCbl-replete environments (18).

**Table 1.**
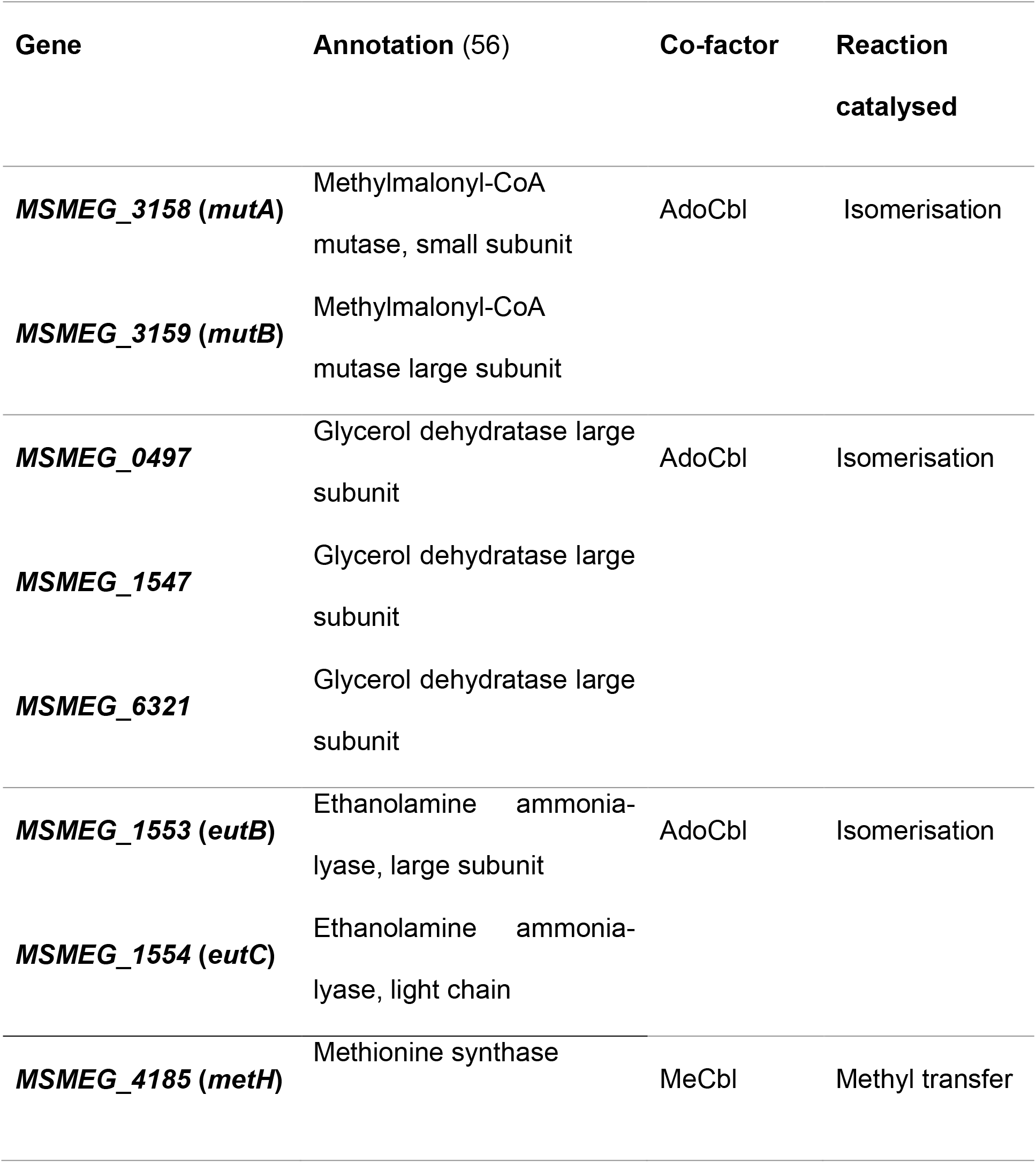
Predicted cobalamin-dependent enzymes in *M. smegmatis*.

We postulated that *metE* would similarly be subject to riboswitch-mediated repression in *M. smegmatis*, given the inferred genetic capacity for *de novo* cobalamin biosynthesis in this organism (5). Moreover, the cobalamin-mediated repression of *metE* would render *metH* essential for growth of *M. smegmatis in vitro*. In this study, we provide direct biochemical confirmation that *M. smegmatis* constitutively produces cobalamin *in vitro*. We further show that *M. smegmatis* utilises exogenous CNCbl and dicyanocobinamide ((CN)_2_Cbi) as precursors for the biosynthesis of the physiologically relevant cobalamin co-factor. However, our results indicate that the uptake of these corrinoid precursors by *M. smegmatis* is restricted. Finally, we demonstrate that the expression of *metE* in *M. smegmatis* is under constitutive repression by a cobalamin riboswitch, a finding which explains the essentiality of *metH* in this non-pathogenic mycobacterium under standard culture conditions.

## RESULTS

### *The de novo* cobalamin biosynthesis pathway is functional in *M. smegmatis*

Genomic analyses indicate that *M. smegmatis* encodes the complete pathway for *de novo* cobalamin biosynthesis (5). To investigate the ability of *M. smegmatis* to synthesise cobalamin using the *de novo* pathway, we developed a liquid chromatography tandem mass spectrometry (LC-MS/MS) method based on the derivatisation of cobalamin to CNCbl by potassium cyanide (KCN). Then, utilizing multiple reaction monitoring (MRM) of two co-eluting transitions corresponding to α-ribazole 5’-phosphate and DMB (Figure 1A), we identified cobalamin in the cell extracts as derivatised CNCbl, validated by two transitions co-eluting at 2.79 min (Figure 1C). Using this method, high intensity CNCbl peaks were identified in cell extracts of wild type *M. smegmatis* mc^2^155 grown aerobically to stationary phase in standard Middlebrook 7H9-OADC medium (Figure 1D). To confirm *de novo* cobalamin production in *M. smegmatis*, we generated an unmarked, in-frame deletion of *cobK* (Figure S1A, C-E), which encodes a putative precorrin-6x reductase required for corrin ring synthesis. To ensure that all phenotypes reflected the consequences of the specific gene deletions and/or disruptions, and were not confounded by off-site mutations, the parental strain and derivative mutants were subjected to whole-genome sequencing. Single nucleotide mutations (SNMs) in six genes were uniquely identified in the Δ*cobK* strain but not in the parental wild type strain but none of the SNMs could be linked to cobalamin or methionine metabolic pathways (Table 2).

**Table 2.** SNMs unique to the Δ *cobK* strain relative to the wild type parental strain.

| Gene | Annotation (56) | SNP | Type of SNM | Amino Acid Change |
| --- | --- | --- | --- | --- |
| **MSMEG_0691** | Putative transcriptional regulatory protein | 1T>G | 5' UTR mutation |  |
| **MSMEG_2148** | HNH endonuclease domain protein | 1135 T>C | missense | Trp379Arg |
| **MSMEG_3876** | Putative phosphotransferase enzyme family protein | 887G >A | missense | Arg296His |
| **MSMEG_6127** | Anti-anti-sigma factor | 107T >G | missense | Leu36Arg |
| **MSMEG_6270** | Hypothetical protein | 1A>C<br>1G>A | 5' UTR mutation |  |
| **MSMEG_6423** | Glycerophosphoryl diester phosphodiesterase family protein | 1T>C<br>1T>G | 5' UTR mutation |  |

In contrast to wild type cell extracts, the Δ*cobK* extracts lacked the dual co-eluting peaks characteristic of CNCbl (Figure 1E). This observation indicated the indispensability of CobK for cobalamin biosynthesis and provided further evidence that the cobalamin signal detected in the wild type strain (Figure 1D) resulted from *de novo* biosynthesis. We also confirmed the absence of cobalamin production in a double Δ*metE cobK*::*hyg* knock-out (KO) strain during growth in L-methionine-supplemented medium (Figure 1F). This strain, in which the entire *metE* open reading frame (ORF) is deleted and *cobK* is disrupted by the insertion of a hygromycin (*hyg*) resistance marker (Figure S1B, S1F), is a methionine auxotroph that can only be propagated in media supplemented with methionine or CNCbl (Figure 1G).

### *M. smegmatis* assimilates exogenous CNCbl and (CN)_2_Cbi *in vitro*

The ability to propagate the Δ*metE cobK*::*hyg* mutant in media containing methionine or CNCbl indicated that *M. smegmatis* can utilise exogenous methionine and CNCbl in the absence of an intact *de novo* cobalamin biosynthesis pathway, pointing to functional transport and assimilation pathways. We previously showed that *M. tuberculosis* could utilise dicyanocobinamide ((CN)_2_Cbi) during growth *in vitro* (12). To determine whether *M. smegmatis* could also assimilate this cobalamin precursor, we tested the ability of the Δ*metE cobK*::*hyg* double mutant to grow in media supplemented with (CN)_2_Cbi. First, the strain was grown to exponential phase with excess methionine (1mM), after which 10-fold serial dilutions were spotted onto Middlebrook 7H10-OADC agar containing 10μM (CN)_2_Cbi. After 3 days’ incubation at 37°C, (CN)_2_Cbi uptake was qualitatively assessed by examining colony sizes (Figure 2A). For comparison, serial dilutions were also spotted on agar supplemented with 10μM CNCbl. Interestingly, the growth of Δ*metE cobK*::*hyg* strain was very limited on agar supplemented with (CN)_2_Cbi (Figure 2A). In fact, growth was observed only in the most concentrated (undiluted) bacterial spots (Figure 2A). While this might have been as a consequence of methionine carryover, the fact that similar growth was not observed in the unsupplemented 7H10 plate (Figure 2A) suggested that was not the case. Instead, these observations implied the ability of *M. smegmatis* to utilise (CN)_2_Cbi, albeit to a much lesser extent than CNCbl (Figure 2A). To test the potential for (CN)_2_Cbi to support growth in liquid culture, an “MIC-type” Alamar Blue assay (20) was performed in which growth from an inoculum of ~5 × 10^3^ Δ*metE cobK*::*hyg* cells was determined in media containing 2-fold serial dilutions of (CN)_2_Cbi (Figure 2B). The Δ*metE cobK*::*hyg* strain was viable at (CN)_2_Cbi concentrations higher than 7.5μM (Figure 2B), consistent with the ability to convert the corrinoid precursor to cobalamin.

**Figure 2.**
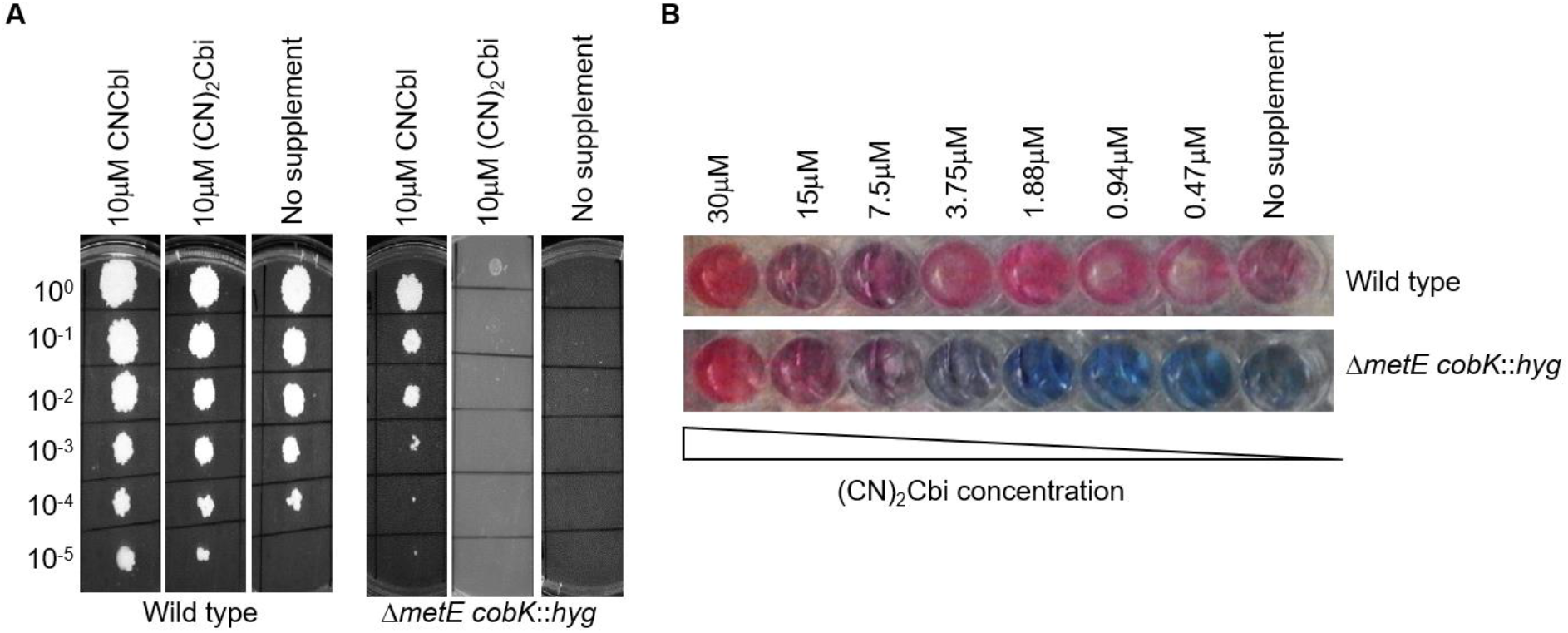
Uptake of exogenous CNCbl and (CN)_2_Cbi in *M. smegmatis*. **A**. Spotting assays of exponential-phase cultures of wild type and Δ*metE cobK*::*hyg* strains on 7H10-OADC agar with or without 10μM CNCbl or 10μM (CN)_2_Cbi shows restricted uptake of (CN)_2_Cbi relative to CNCbl uptake on solid medium. **B**. Alamar Blue assay to evaluate the growth of the Δ*metE cobK*::*hyg* strain in liquid medium supplemented with (CN)_2_Cbi. 5 × 10^3^ cells were seeded in 7H9-OADC medium supplemented with 2-fold dilutions of (CN)_2_Cbi starting at 30μM as the highest concentration.

To confirm the assimilation of (CN)_2_Cbi in *M. smegmatis*, we used LC-MS/MS to analyse cell extracts of wild type, Δ*cobK* and Δ*metE cobK*::*hyg* strains grown to stationary-phase in 7H9-OADC medium with or without excess (30μM) (CN)_2_Cbi (Figure 3A-F). As a positive control for *de novo* cobalamin biosynthesis, we analysed the wild type strain grown in parallel without supplementation (Figure 3B). For the Δ*metE cobK*::*hyg* mutant, methionine supplementation was used to enable propagation in the absence of (CN)_2_Cbi (Figure 3C). We observed that all the (CN)_2_Cbi-supplemented strains reached stationary phase simultaneously. However, the dual co-eluting peaks characteristic of CNCbl were detected only in the (CN)_2_Cbi-supplemented Δ*metE cobK*::*hyg* strain (Figure 3C, D), strongly suggesting uptake and conversion of (CN)_2_Cbi to cobalamin. Interestingly, cobalamin was not detectable in the (CN)_2_Cbi-supplemented Δ*cobK* strain (Figure 3E, F), which did not require supplementation for growth. The assimilation of (CN)_2_Cbi in the Δ*metE cobK*::*hyg* strain was also accompanied by a distinct change in the colour of the spent media from purple to pale yellow (Figure 3C, inset). By comparison, the colour of the spent media in the (CN)_2_Cbi-supplemented wild type and Δ*cobK* cultures changed only slightly to a rusty hue (Figure 3B (inset) & Figure 3F (inset)), consistent with limited uptake in these strains.

**Figure 3.**
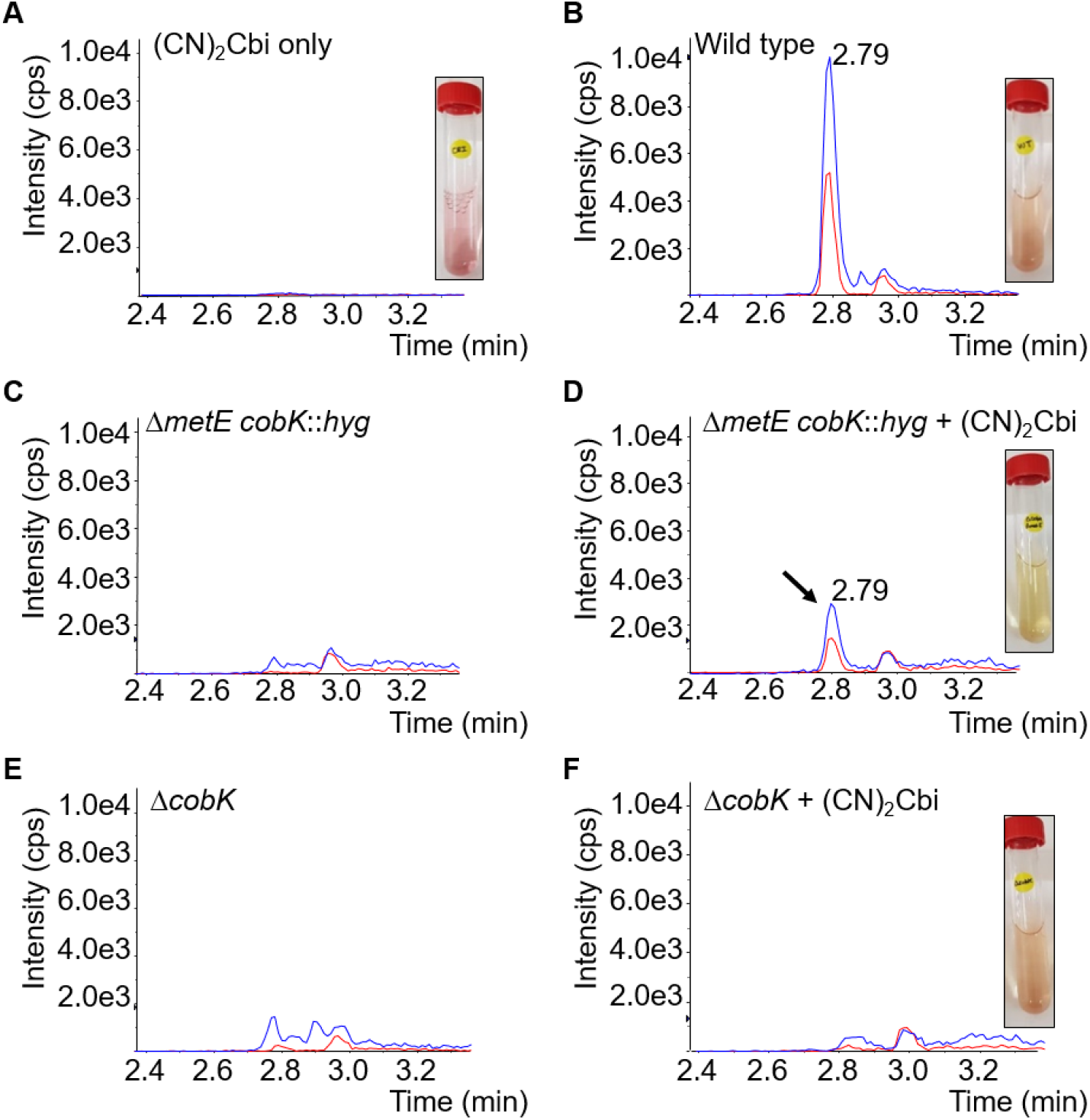
Assimilation of (CN)_2_Cbi in *M. smegmatis*. **A**. (CN)_2_Cbi control. **B**. *De novo*-synthesised cobalamin in the wild type strain. **C-D**. Detection of recovered cobalamin due to (CN)_2_Cbi assimilation in the Δ*metE cobK*::*hyg* double mutant. (**E-F**). Absence of recovered cobalamin in the Δ*cobK* strain in the presence of exogenous (CN)_2_Cbi. (CN)_2_Cbi uptake was accompanied by changes in the colour of the spent media from purple (***A, inset***) to a rusty hue in the wild type (***B, inset***) and Δ*cobK* strains (***F, inset***), and pale yellow in the Δ*metE cobK*::*hyg* strain (***D, inset***).

### MetE expression in *M. smegmatis* is regulated by a cobalamin-sensing riboswitch

We previously reported that the cobalamin-sensing riboswitch located in the 5’ UTR of *metE* strongly attenuated the transcription of this gene in *M. tuberculosis* in the presence of exogenous CNCbl (18). To investigate whether the corresponding riboswitch in *M*. *smegmatis* operated similarly, we analysed relative *metE* transcript levels in wild type and Δ*cobK* strains grown in the presence or absence of CNCbl using droplet digital (dd)PCR. We found low but detectable levels of *metE* transcripts in the wild type strain (Figure 4A). By comparison, *metE* transcripts were 19× more abundant in the Δ*cobK* strain (Figure 4A), supporting the notion that the abrogation of *de novo* cobalamin biosynthesis in the mutant released *metE* transcription from riboswitch-mediated repression. There was no significant difference in *metE* transcript levels between the CNCbl-supplemented and unsupplemented wild type strain (Figure 4A). In contrast, a small but statistically significant reduction (0.86×) in *metE* transcript levels was observed in the Δ*cobK* strain in the presence of exogenous CNCbl (Figure 4A). The absent to modest change in *metE* levels in these strains following CNCbl supplementation suggested that the uptake/assimilation of exogenous CNCbl might be restricted in *M. smegmatis*, consistent with the LC-MS/MS results. Alternatively, these results could indicate selective repression of cobalamin uptake/assimilation systems in strains which do not require the co-factor for viability or growth.

**Figure 4.**
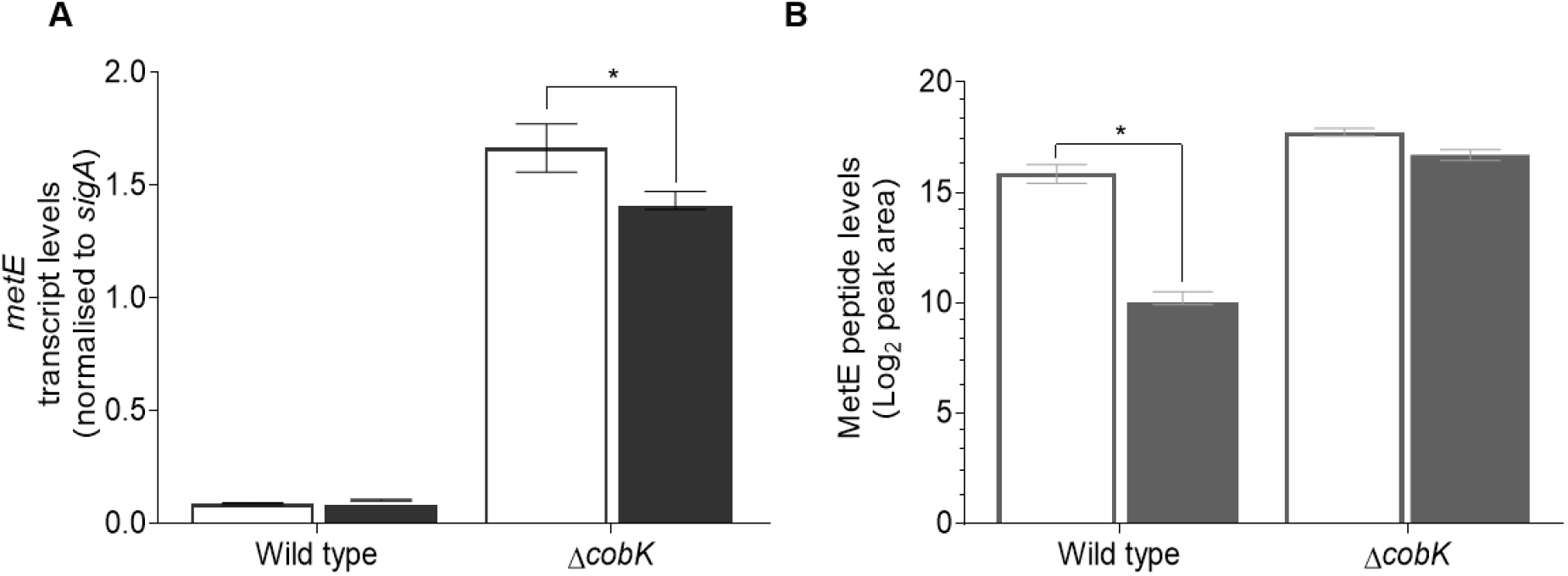
Cobalamin-mediated attenuation of MetE expression in *M. smegmatis*. **A**. ddPCR analysis of *metE* transcription in the wild type and Δ*cobK* strains cultured in the presence (solid bars) or absence (open bars) of exogenous CNCbl. The Δ*cobK* strain exhibited an overabundance of *metE* transcripts relative to wild type. A small but statistically significant decrease in the level of *metE* transcript in the Δ*cobK* strain was observed in the presence of exogenous CNCbl (*p*=0.0359; (*); two-way ANOVA), but the change in *metE* transcript levels in the wild type strain was not statistically significant. The graphed data are representative of two independent experiments. Error bars show the standard error of the mean. **B**. Targeted MS analysis of MetE peptide levels (log_2_ peak area) in the wild type and Δ*cobK* strains grown in the presence (solid bars) or absence (open bars) of exogenous CNCbl. Exogenous CNCbl more significantly decreased MetE peptide levels in the wild type strain relative to the Δ*cobK* mutant (*p*=0.0151; (*) two-way ANOVA).

To examine how cobalamin availability in *M. smegmatis* affected MetE protein content, we adapted a targeted MS method (21) to measure MetE peptide levels in wild type and Δ*cobK* strains grown in the presence or absence of exogenous CNCbl. This analysis indicated that MetE protein levels were 3× more abundant in the Δ*cobK* mutant than in the parental wild type strain (Figure 4B). In the presence of exogenous CNCbl, the Δ*cobK* mutant exhibited a 2× decrease in MetE protein levels (Figure 4B). Unexpectedly, exposure of the wild type strain to exogenous CNCbl caused a 48× reduction in MetE protein levels (Figure 4B). This result contrasted with the modest impact of CNCbl on *metE* transcript levels (Figure 4A) and suggested that MetE expression was likely primarily controlled at the translational level by this riboswitch.

### *metH* is essential in *M. smegmatis*

The inferred cobalamin-mediated repression of *metE* in turn implied that MetH function would be indispensable for the growth of wild type *M. smegmatis*. To test this prediction, we attempted to generate an in-frame *metH* deletion mutant (Figure S1A) by two-step allelic exchange mutagenesis (22). The Δ*metH* construct was designed to mimic a naturally-occurring *metH* truncation which partially disrupts the cobalamin-binding domain and eliminates the *S*-adenosyl-L-methionine (SAM)-binding domain of MetH in *M. tuberculosis* CDC1551 (18). Another *metH* KO construct containing a *hyg* marker (Figure S1A) was also designed to enable the recovery of *metH* double crossover (DCO) mutants by “forced” selection on Hyg. Anticipating the loss of viability owing to *metH* essentiality, all media were supplemented with 1mM L-methionine. Of 154 putative *metH* DCO recombinants screened by PCR, none (0/154) carried the Δ*metH* allele; instead, all 154 colonies were wild type revertants. Similarly, 60 putative *hyg*-marked DCOs were screened by PCR, none of which bore the Δ*metH* allele. These results strongly suggested that *metH* was essential in *M. smegmatis*, consistent with recent genetic screens which identified *metH* among the subset of essential genes in *M. smegmatis* (23, 24).

### Conditional depletion by CRISPRi confirms *metH* essentiality

Since our attempts to delete *metH* in *M. smegmatis* were unsuccessful, we instead opted to generate a *metH* conditional knock-down (cKD). For this purpose, we employed the anhydrotetracycline (ATc)-inducible mycobacterial CRISPRi system (25), utilizing a panel of 13 short guide (sg)RNAs targeting different regions of the *metH* ORF and with different target complementarity scores (Table S3). An sgRNA targeting *mmpL3*, an essential gene involved in mycolic acid biosynthesis (26), was used as a positive control. Gene silencing in transformed cells was assessed by growth inhibition on ATc-containing selection media. The induction of *metH* silencing by ATc inhibited colony formation in wild type *M. smegmatis*, confirming the essentiality of *metH* under the conditions tested (Figure 5A). Consistent with previous work (24, 25), sgRNAs with higher complementarity scores displayed stronger gene silencing, leading to more robust growth inhibition than sgRNAs with lower scores (Figure S2). ATc-dependent growth inhibition was rescued by supplementation with exogenous methionine (Figure 5A), indicating that the lack of growth in the *metH* cKD strains resulted from methionine starvation.

**Figure 5.**
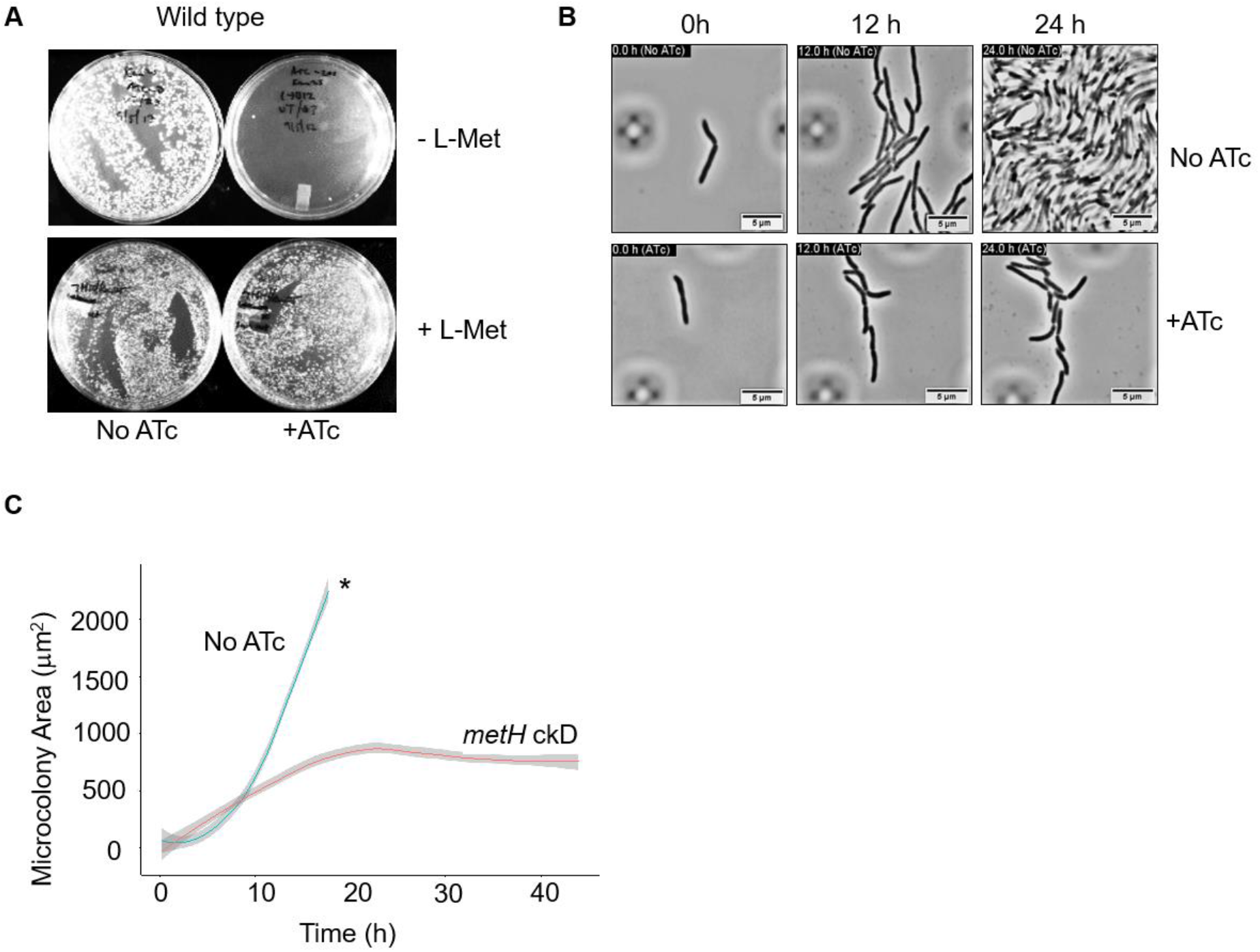
Growth cessation due to methionine depletion in the *metH* cKD. **A**. ATc-induced growth inhibition in the *metH* cKD is rescued by exogenous methionine (L-Met). **B**. Representative images from time-lapse microscopy at the 0-h, 12-h, and 24-h time points showing severe growth retardation in the *metH* cKD. Images are taken from Supplementary Movie 2. Scale bars, 5μm. **C**. Quantification of microcolony growth in the *M. smegmatis metH* cKD using “R” software. * limit of detection.

To investigate this phenotype further, we traced the growth of the *metH* cKD strain at the single-cell level using microfluidics and time-lapse microscopy. A log-phase culture of cells carrying the *metH* cKD construct was pre-incubated with ATc for 6 h at 37°C and then loaded into the CellASIC® ONIX2 microfluidic device and imaged in real time over the course of 43 h with constant perfusion with 7H9-OADC media containing ATc. In parallel, an uninduced (No ATc) control was perfused with 7H9-OADC media only. Analysis of the time-lapse images showed that the induced *metH* cKD strain shared similar morphological features of shape and size with the No ATc control (Figure 5B). Moreover, in both the *metH* cKD strain and the No ATc control, cells divided by v-snapping (Supplementary Movies 1-5), which is typical of mycobacterial cell division (27, 28). However, the growth rates of the *metH* cKD strain and the No ATc control were markedly different (Figure 5C). In the No ATc control, microcolonies displayed an exponential-phase doubling time of 2.94 ± 0.3 h (Figure 5C). In this control, tracing of distinct cells was feasible only for the first 18 h of the experiment; by 24 h, microcolonies had attained confluence, occupying the entire field of view (Figure 5B; Supplementary Movies 1-5). Consistent with methionine depletion in the *metH* cKD strain, this mutant exhibited a much slower replication rate, doubling every 5.23 ± 0.4 h until the 18-h time-point when the growth rate flat-lined (Figure 5C) and the expansion of microcolonies slowed and appeared to halt by 24 h (Figure 5B; Supplementary Movies 1-5).

### Abrogation of cobalamin biosynthetic capacity alleviates *metH* essentiality in *M. smegmatis*

The time-lapse microscopy data suggested that the essentiality of the *metH* gene might depend on both the presence of endogenous cobalamin and a functional *metE* riboswitch in *M. smegmatis*. Therefore, we reasoned that it would be possible to create a *metH* deletion in the cobalamin-deficient Δ*cobK* strain. To test this hypothesis, we generated an unmarked in-frame *metH* deletion in the Δ*cobK* background and screened the resultant DCOs by PCR. As expected, PCR screening identified 9 out of 34 putative DCOs as Δ*cobK* Δ*metH* double KO mutants (Figure S1C-E), linking the essentiality of *metH* to endogenous cobalamin availability. We found that silencing of *metH* had no effect on the viability of the cobalamin-deficient Δ*cobK* mutant (Figure 6A), confirming that endogenous cobalamin was required to block methionine biosynthesis via riboswitch-mediated repression of *metE*. Moreover, consistent with the limited impact of exogenous CNCbl on MetE protein levels (Figure 4B), CNCbl supplementation had a negligible effect during the growth of the Δ*cobK* strain on solid media following ATc-induced *metH* silencing (Figure 6A).

**Figure 6.**
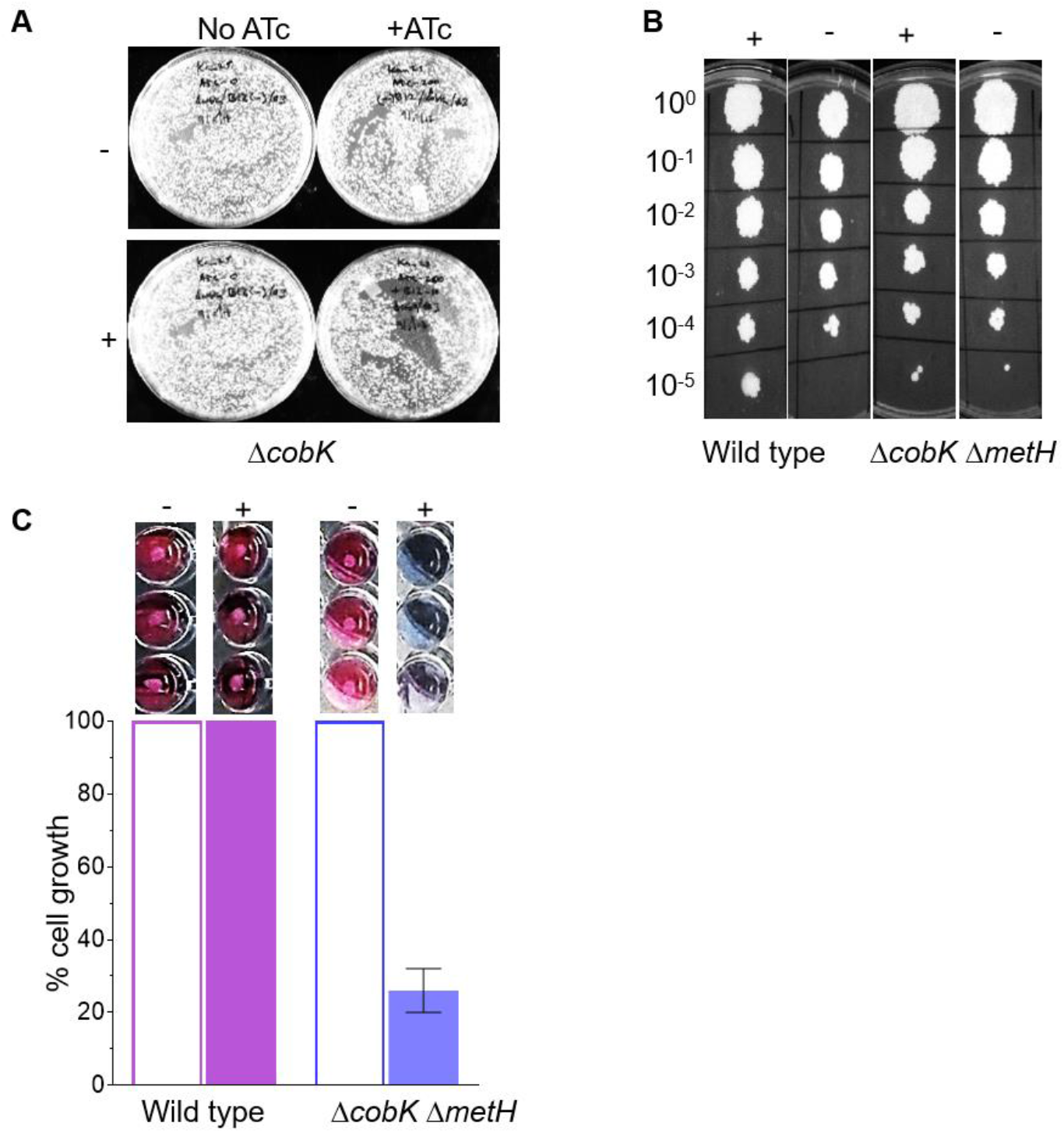
Sensitivity of MetH mutants to exogenous CNCbl. **A**. CRISPRi-mediated silencing of *metH* in the Δ*cobK* strain in the presence (+) or absence (-) of exogenous CNCbl. Exogenous CNCbl had negligible effect on the growth of *metH* cKD on solid media in this background. **B.** A spotting assay on 7H10 agar containing 10μM exogenous CNCbl showing the failure of exogenous CNCbl to inhibit growth of the Δ*cobK* Δ*metH* strain on solid media. **C**. Sensitivity of the Δ*cobK* Δ*metH* strain to exogenous CNCbl in liquid medium as determined using the Alamar Blue assay.

Next, we investigated the impact of exogenous CNCbl during growth of the Δ*cobK* Δ*metH* strain on solid *versus* in liquid media. To this end, a late log-phase (OD_600_ ~ 1) culture was 10-fold serially diluted and spotted on 7H10-OADC agar supplemented with or without 10μM CNCbl (Figure 6B). There was no impairment of growth of the CNCbl-supplemented Δ*cobK* Δ*metH* mutant on solid media (Figure 6B). To determine if this phenotype were also true in liquid media, we seeded an inoculum of 2.5 × 10^3^ Δ*cobK* Δ*metH* cells and analysed cell proliferation after an overnight incubation at 37°C with or without 10μM CNCbl using the Alamar blue assay (20) (Figure 6C). Interestingly, CNCbl supplementation led to approximately 80% inhibition of the growth of the Δ*cobK* Δ*metH* strain (Figure 6C). This result confirmed the ability of *M. smegmatis* to assimilate exogenous CNCbl, although the uptake of the corrinoid was seemingly better in liquid than on solid media.

## DISCUSSION

The production of cobalamin by *M. smegmatis* was previously inferred indirectly from microbiological assays (29–31). Using targeted LC-MS/MS approach, we provide direct proof of constitutive *de novo* cobalamin biosynthesis in *M. smegmatis* under aerobic conditions. The LC-MS/MS method optimised in this work utilised MRM of two mass spectra corresponding to DMB, the lower base in physiologically relevant cobalamin. Hence, we infer that *M. smegmatis* is able to synthesise and use DMB. Unconventional cobamides known as “pseudo-coenzyme B_12_” comprising an α-axial ligand other than DMB, but with the adenosine group retained, have been found in other bacteria (32). *M. smegmatis* encodes a CobT protein, which is closely homologous to those of *S. meliloti* and *S. enterica*, both of which incorporate a range of α-ligands in their cobamide structures (33). While there has been no evidence that mycobacteria produce pseudo-coenzyme B_12_, it is possible that our DMB-dependent detection method did not capture the full range of cobamides present in *M. smegmatis*. Nonetheless, these results indicated substantial cobalamin production in *M. smegmatis*, consistent with an early study which detected low-level cobalamin production in *M. smegmatis* using the *L. leichmannii* tube assay (29). The disruption of *cobK*, encoding a predicted precorrin-6x reductase, abrogated cobalamin production, confirming that a single genetic lesion can cripple the entire pathway. It is noteworthy, therefore, that the loss of *cobF*, encoding a putative precorrin-6a synthase occurring immediately upstream of CobK in the pathway, has been identified as one of the defining molecular events in the evolution of pathogenic mycobacteria (34–36).

Both CNCbl and (CN)_2_Cbi supported the growth of the Δ*metE cobK*::*hyg* strain by enabling MetH-dependent production of methionine. These results demonstrate that *M. smegmatis* is capable of corrinoid transport and assimilation. Since CNCbl and (CN)_2_Cbi must undergo decyanation and adenosylation to produce cobalamin, *M. smegmatis* also encodes as yet uncharacterised enzymes for these reactions. Interestingly, our data showed a seemingly enhanced capacity for (CN)_2_Cbi assimilation in the Δ*metE cobK*::*hyg* strain compared to wild type and Δ*cobK* strains. Since both these strains are able to grow without cobalamin owing to the presence of MetE as alternative methionine synthase, the reduced (CN)_2_Cbi uptake might suggest the potential for selective assimilation/transport as a function of methionine biosynthetic capacity. The peak intensity of the recovered cobalamin in the (CN)_2_Cbi-supplemented Δ*metE cobK*::*hyg* strain was significantly lower than that of *de novo*-synthesised cobalamin in the wild type (Figure 3B, D), possibly indicating that much lower amounts of co-factor are necessary to support growth.

In *M. tuberculosis,* the nonspecific ABC-type transporter, BacA (Rv1819c) has been identified as the sole cobalamin and corrinoid transporter (12). Until recently, the mechanistic details of cobalamin transport by Rv1819c had remained elusive. However, the resolution of the crystal structure of Rv1819c (37) has provided key insights into its function in the uptake of hydrophilic molecules, suggesting that this protein passes a cargo slowly along its cavity via facilitated diffusion. Facilitated diffusion is a very low-efficiency process and, if the *M. smegmatis* homologue functions similarly in corrinoid uptake, it might explain the poor uptake of CNCbl and (CN)_2_Cbi. *M. smegmatis* also contains two predicted Rv1819c homologues, encoded by paralogous genes located at different genomic loci (MSMEG_3655 & MSMEG 4380). In addition, *M. smegmatis* encodes an operon containing putative homologues of BtuF (MSMEG_4560), BtuC (MSMEG_4559) and BtuD (MSMEG_4558), all components of the classic TonB-ExBD-BtuFCD cobalamin transport system in Gram-negative bacteria (38). Whether these genes encode functional transporters is still unknown and further research is needed to determine which proteins are involved in ferrying corrinoids and their precursors across the notoriously complex mycobacterial cell wall (39).

We previously reported that a cobalamin-sensing riboswitch controlled *metE* transcription in *M. tuberculosis* (18). In that work, the level of *metE* transcript was decreased in the presence of exogenous CNCbl, leading to the conclusion that this riboswitch functioned as a transcriptional “off” switch. In the current study, we found that the levels of *metE* transcript were much lower in the cobalamin-replete wild type *M. smegmatis* strain compared to the Δ*cobK* mutant. Since riboswitches sense ligand levels to attenuate expression (40), the low-level *metE* transcripts found in the wild type strain likely reflects a physiological equilibrium between ligand-bound and unbound riboswitch states which allow for limited gene expression. Therefore, although the uptake of exogenous CNCbl is restricted in both wild type and Δ*cobK* strains, the low level of uptake was still enough to shift the endogenous ligand-riboswitch equilibrium more significantly in wild type than in mutant cells which exhibited elevated MetE protein content (Figure 4A, B). Unlike in *M. tuberculosis*, exogenous CNCbl was unable to exert significant changes to *metE* transcript levels in *M. smegmatis*, presumably owing to the limited uptake. While these results imply transcriptional regulation, we also observed an unexpected and dramatic reduction in MetE protein levels in the wild type in the presence of exogenous CNCbl. These results suggested that this riboswitch might utilise a coupled translational-transcriptional regulation mechanism by which the inhibition of translation initiation precedes transcription termination and mRNA instability (41–43). Future work will elucidate the precise mechanism of cobalamin-sensing riboswitches in mycobacteria.

Our *in vitro* results support the conclusion that constitutive endogenous production of cobalamin compels *M. smegmatis* to rely on MetH for the biosynthesis of methionine. We found that the disruption of MetH activity was growth-retarding in the presence of cobalamin, ostensibly owing to methionine depletion. In contrast, in the absence of cobalamin, bacilli were relieved of riboswitch-mediated repression of MetE, allowing the alternative methionine synthase to substitute for the inactivated MetH. We predict that *metH* will be essential in all mycobacteria capable of *de novo* cobalamin biosynthesis, representing an important deviation from cobalamin-deficient pathogenic mycobacteria like *M. tuberculosis*. The corollary is that mycobacterial species which are incapable of *de novo* cobalamin biosynthesis will accommodate MetH inactivation. Indeed, *metH*-null *M. tuberculosis* mutants have been generated (18) and several naturally-occurring, potentially inactivating mutations in *metH* have been found in circulating *M. tuberculosis* clinical isolates (13). Consistent with our findings, a recent Tn-screen identified *metH* among the subset of genes essential for the growth of *M. smegmatis in vitro* (23). Another recent study reported the inactivation of MetH in *M. smegmatis*, but in contrast to our findings, the authors did not observe any growth inhibition in their *metH*-null mutants in supplement-free media (31). The results presented here, together with our own independent Tn-seq and CRISPRi-seq analyses of *M. smegmatis* gene essentiality (24), demonstrate that *metH* cannot be disrupted in a cobalamin-replete strain without sacrificing viability. This apparent conflict might be explained by the possibility that the *metE* riboswitch in the parental strains used by Guzzo *et al.* to generate their *metH*-null mutants harboured inactivating mutations, which can accumulate spontaneously during the serial passage of mycobacterial cultures (44, 45). To eliminate the potential confounding effect of mutations in our study, the wild type and its derivative mutant strains were subjected to whole-genome sequencing.

In summary, we have shown that *M. smegmatis*, a non-pathogenic mycobacterium, is a constitutive producer of cobalamin *in vitro*. Surprisingly, the transport of corrinoids in *M. smegmatis* appears restricted despite the presence in the genome of multiple putative transporters. Notably, this study also revealed differences in the regulation of methionine biosynthesis between *M. smegmatis* and *M. tuberculosis*. These differences in cobalamin-dependent metabolism between an environmental mycobacterium and an obligate pathogen might be informative in understanding the selective pressures which have shaped *M. tuberculosis* metabolism for pathogenicity and host tropism.

## MATERIALS AND METHODS

### Bacterial strains and culture conditions

The bacterial strains and plasmids used in this study are described in Table S1. Unless specified, *M. smegmatis* cultures were grown in either Middlebrook (Difco) 7H9 broth supplemented with 10% Oleic Albumin Dextrose Catalase (OADC) (Becton Dickinson) and 0.05% Tween 80 or on Middlebrook (Difco) 7H10 agar supplemented with 10% OADC. For mycobacterial cultures, kanamycin (Km) and hygromycin (Hyg) were used at final concentrations of 25μg/mL and 50μg/mL, respectively. *E. coli* was cultured in LB or LA with 50μg/mL Km or 200μg/mL Hyg, where appropriate. All cultures were incubated at 37°C. To generate growth curves, 50μL *M. smegmatis* cells were seeded at a concentration of 1 × 10^6^ cfu/mL in 96-well culture plates (Greiner Bio-One) and absorbance measurements were recorded every 1.5 h, over a period of 30 h, in a FLUOstar OPTIMA microplate reader (BMG Labtech).

### Cloning

The oligonucleotides (oligos) used for cloning and PCR are listed in Table S2. An in-frame, unmarked deletion in *M. smegmatis cobK* (*MSMEG_3875*) was generated by joining a 912-bp PCR-generated fragment (FR1) containing 40bp of the 5’ end of *cobK* to a second 923-bp PCR-generated fragment (FR2) containing 107bp of the 3’ end of *cobK* in a three-way ligation reaction with p2NIL backbone (Addgene plasmid #20188; (46)), using *Asp*718I, *Bgl*II, and *Hin*dIII restriction. The resultant vector (p3875K) contained a deleted 120-bp *cobK* allele. To generate an in-frame, unmarked deletion in *M. smegmatis metH* (*MSMEG_4185*), an 1524-bp amplicon (FR1) of the 5’ coding sequence of *metH* and another 1480-bp amplicon (FR2) containing 354bp of the 3’ end of *metH* were joined in a three-way ligation reaction with p2NIL using *Asp*718I, *Hin*dIII and *Bgl*II to produce the p4185K vector carrying a truncated *metH* allele of 1848bp. Counter-selection fragments carrying the *lacZ*, *hyg*, and *sacB* genes was excised from pGOAL19 (Addgene plasmid #20190; (46)) and cloned at *Pac*I sites of p3875K and p4185K to generate the suicide vectors p3875K19 and p4185K19, respectively. To generate the *hyg*-marked *metH* construct, a *hyg* cassette was excised from the pIJ963 vector (47) and cloned into the *Bgl*II site of p4185K. A counter-selection cassette derived from pGOAL17 (Addgene plasmid #20189; (46)) was then inserted into p4185K to generate p4185K17. Constructs were validated by restriction enzyme mapping and Sanger sequencing using the primers listed in Table S2.

### Isolation of allelic exchange mutants

*M. smegmatis* Δ*cobK* and Δ*cobK* Δ*metH* mutants were generated by allelic exchange mutagenesis (22). A total of 100μL competent cells were incubated in a 1mm cuvette with 1–8μg DNA for 20 min on ice prior to pulsing in a GenePulser Xcell™ electroporator (Bio-Rad) with time constant and voltage settings at 5 ms and 1200V, respectively. Single crossover (SCO) transformants were selected with Km and Hyg on 7H10-OADC plates. As colonies became visible, 50μL of 2% (w/v) 5-bromo-4-chloro-3-indolyl-β-D-galactoside (X-gal) was underlain in each plate for blue/white screening of SCOs. PCR-verified SCOs were then cultured in antibiotic-free 7H9-OADC, followed by 10-fold serial dilutions and plating on 7H10-OADC containing 2% (w/v) sucrose. DCOs were screened by PCR and confirmed with Southern blotting or Sanger sequencing. For Southern blotting confirmation of Δ*cobK*, 2-3μg DNA was digested overnight with *Sty*I, separated on 1% agarose gel at 80V and transferred and fixed onto a HydrobondTM N+ membrane (Amersham), and hybridised overnight at 42°C with target-specific PCR-generated probes labelled with the ECL Direct Nucleic Acid Labelling and Detection Systems (Amersham). The target DNA fragments were visualised on Kodak hypersensitive X-ray films.

### Cobalamin extraction

Wild type or mutant *M. smegmatis* strains were cultured until stationary phase (OD_600_ ~2) in 50mL 7H9-OADC supplemented with 3μg/mL cobalt chloride. Cells were harvested by centrifugation at 4000 × *g* for 10 min at 4°C, re-suspended in 8mL of 50mM sodium acetate buffer, pH4.5, and stored at −80°C until needed. Once thawed, the cells were lysed by 5 min of sonication using a microtip sonicator set at 30 amplitude,15 s pulse on and 15 s pulse off. Next, 16μL of 100mM KCN was added to the lysed cells and, with the extraction tube tightly closed, the samples were incubated at room temperature for 30 min in a chemical fume hood, followed by boiling at 90°C for 45 min inside the hood. The tube was then cooled on ice briefly and centrifuged at 4°C at 4000 × *g* for 10 min. The supernatant was filtered through a 0.22μm filter and loaded onto a Sep-Pak C18 Plus Light Cartridge (Waters), which had been washed with 5mL 75% (v/v) ethanol and conditioned with 10mL sterile water. Next, the cartridge was washed with 10mL of water and eluted with 75% ethanol, collecting about 15 drops. The eluent was analysed immediately by LC-MS/MS or stored in −20°C until needed. When analysis was done on frozen samples, a centrifugation at 14000 × *g* for 10 min on a bench-top centrifuge was first performed, followed by chloroform purification.

### LC-MS/MS detection and analysis of cobalamin

Eluents were analysed using LC-MS/MS in a positive ionisation mode and quantitated using the following multiple reaction monitoring (MRM) parameters: m/z, 678→359 and m/z, 678→147. Chromatographic separation was performed through a HPLC reverse phase column (Phenomenex SynergiTM Polar-RP 100Å, 50 x 2mm (Separations)) using an Agilent 1200 Rapid Resolution HPLC system equipped with a binary pump, degasser, and auto sampler, coupled to an AB Sciex 4000 QTRAP hybrid triple quadrupole linear ion-trap spectrometer. Mobile phases were A: 0.1% formic acid in water; and B: 0.1% formic acid in acetonitrile. The following gradients were run: 0-2 min, 95% A; 2-4 min, 5% A; 4-6 min, 95% A; and 6-8 min, 95% A at a flow rate of 400μL/min. The mass spectrometry analysis was performed on an AB Sciex 4000 QTRAP LC mass spectrometer using the following parameters: Curtain gas (25.00); IS (5500.00); Temperature (200.00°C); GS1 (80.00); GS2 (55.00); EP (12.0). Data processing was done using the SCIEX Analyst® software.

### Quantitative gene expression analysis by ddPCR

Droplet digital PCR (ddPCR) and data analysis was performed as described previously (48). Total RNA was extracted using the FastRNA® Pro Blue kit (MP Biomedicals) and DNase-treated with TURBO DNAse (Ambion) after which 0.5μg was used as template for cDNA synthesis, using the High Capacity RNA to cDNA kit (Thermofisher). Primers and minor groove binder (MGB) Taqman probes (Table S2) were designed using Primer Express 3.0 (Applied Biosystems). For duplexing, TaqMan MGB probes homologous to the target genes were labelled with 2’-chloro-7’-phenyl-1, 4-dichloro-6-carboxyfluorescein (VIC) whereas those binding the reference gene, *sigA*, were labelled with 6-carboxyfluorescein (FAM).

### Targeted protein mass spectrometry

Triplicate cultures of *M. smegmatis* were grown to OD_600_ ~1.2 in 7H9-OADC with or without 10μM CNCbl. Cell lysis, fractionation and the generation of tryptic peptides was done as described (21). Selected Reaction Monitoring (SRM) assays were developed in Skyline (version 4.1) using a spectral library generated from previous discovery MS data (21) with a cut-off score of 0.9. Skyline was set up to select two peptides per input protein, with the highest picked MS1 intensity in the discovery data, and then the top 5 most intense fragment ions for each of those peptides. A transition list was then generated for the Thermo Scientific triple stage quadrupole (TSQ) Vantage mass spectrometer. Samples were separated using a Thermo Accella LC system on a 10-cm monolithic C18 column (Phenomenex) with a 4.6-mm ID with a mobile phase that comprised a mixture of solvent A (water + 0.1% formic acid) and solvent B (HPLC-grade acetonitrile + 0.1% formic acid). The method run time was 45 min in total with a flow rate of 300μL/min. The gradient program began with 3% B, followed by a gradient of 8%-45% B from 5-25 min, then an increase to 80% B at the 30-min mark for a 5-min wash, before returning to 3% B for the remainder of the method. The LC system was run in-line into a Thermo TSQ Vantage through a heated electrospray ionisation (HESI) source. The source voltage was +3500V, with a capillary temperature of 300°C, a vaporiser temperature of 200°C, sheath gas of 30, and aux gas of 10. To determine the retention time for each peptide, methods were generated for the TSQ Vantage with a maximum of 20 transitions monitored per method. Since the original list contained 5 transitions, a total of 8 unscheduled methods were generated, with a cycle time of 5 s to maximise the amount of signal, a collision gas pressure of 1.5 mTorr, a Q1 peak width (FWHM) of 0.7, and collision energies as determined by Skyline. The unscheduled methods were then run with consecutive 2μL injections of a reference sample to further refine the list of transitions and determine the retention time for each peptide. The reference sample was generated by pooling all samples. The unscheduled runs were analysed in Skyline to determine the retention times for each peptide. Any transitions with no intensity, background-level intensity, interference, or ambiguous signal were removed from the method, and a minimum of 3 transitions per peptide were kept in the final list. The spectral library was used to further refine the assays and any peptides with a *dotp* score lower than 0.7 were removed from the final list.

### Microplate Alamar Blue Assay

Cell viability was determined using the microplate Alamar blue assay (20) as follows: 50μL of 1:1000 diluted exponential-phase cultures (OD_600_ ~ 0.5) was added to 50μL 7H9-OADC with or without 10μM CNCbl in a 96-well plate. Plates were incubated overnight at 37°C, after which 10μL of 100μg/mL resazurin was added to each well. The plates were incubated for an additional 5 h at 37°C before fluorescence intensity measurements were taken using a FLUOstar OPTIMA microplate reader (BMG Labtech) using excitation and emission wavelengths of 485 nm and 508 nm, respectively.

### Gene silencing using CRISPRi

Thirteen pairs of sgRNA oligos targeting the *M. smegmatis metH* ORF (Table S3) were designed as described previously (24). The oligos were annealed and cloned into the PLJR962 plasmid using *Bsm*BI restriction sites in an overnight ligation reaction with T4 DNA ligase (NEB). Following ligation, the entire reaction mix (10μL) was transformed into 50μL electrocompetent *E. coli* DH5α cells and selected on LB plates with 50μg/mL Km. Plasmid DNA was extracted from single colonies and validated by Sanger sequencing using primer 1834 (Table S3). Next, competent *M. smegmatis* cells were transformed by electroporation with 200ng of *metH* cKD constructs or an *mmpL3* cKD control and selected on 7H10-OADC containing 25μg/mL Km with or without 100ng/mL ATc

### WGS, genome assembly and variant detection

Genomic DNA was extracted as described by van Helden *et al*. (49) from exponential phase cultures of single colonies. Genomic libraries, prepared using the TruSeq Nano DNA (Illumina) sample preparation kit according to the manufacturer’s instructions, were sequenced using a 150-bp paired-end strategy on an Illumina HiSeq 4000 instrument. Trimmomatic v0.35 (50) was used to remove adapters, leading or trailing bases with a quality score < 3, reads shorter than 36bp in length, and bases with an average quality score of < 15 based on a 4-base sliding window. BWA v0.7.12 (51, 52) was then used to map paired-end reads to the *M. smegmatis* mc^2^155 reference genome (CP000480.1) (53). SAMtools v0.1.2 (54) was used to call bases. Sites that had Phred scores lower than 20 or coverage below 10-fold were removed from further analysis. SNPeff v4.1 (55), using the *M. smegmatis* mc^2^155 (uid57701) reference, was used to annotate variant positions.

### Live-cell imaging and quantification of the growth of microcolonies

A 100μL bacterial suspension of 2.0 × 10^6^ cells/mL was prepared and loaded on the four-chambered CellASIC ONIX B04A-03 microfluidic platform (Merck). Cells were trapped with the following pressure and flow time settings: channel A8 at 13.8 kPa for 15 s; channel A6 at 27.6kPa for 15 s. Channel A6 was then rinsed at 6.9kPa for 30 s. Untrapped cells were washed out by flowing inlet solution at 34.5kPa for 5 min. 7H9-OADC medium containing 25μg/mL Km with or without 100ng/mL ATc was perfused continuously for 43 h. Live-cell imaging was performed on a Zeiss AxioObserver using a 100X, 1.4NA Objective with Phase Contrast and Colibri.7 fluorescent illumination system. Images were captured every 15 min using a Zeiss Axiocam 503 and analysed using FIJI software (https://fiji.sc/). To quantify the growth of microcolonies, a thresholded for the time-lapse images was set with a Yen filter in FIJI and the thresholded area over time was then quantified. For each strain type, data extraction and all subsequent analyses were performed on four independent fields of view. The data was analysed using “R” software. Growth curves were generated by subtracting initial background objects from the size data over time and smoothed with a loess regression. Growth rates were predicted from fitting a linear model to the log_2_ microcolony size obtained between 3 h and 18 h.

## ACKNOWLEDGEMENTS

We thank Clemens Hermann and Bridget Calder from Jonathan Blackburn’s proteomics facility for advice and assistance; Sarah Fortune and Jeremy Rock for kindly providing the mycobacterial CRISPRi system; and Stephanie Dawes for providing the Δ*metE cobK*::*hyg* mutant. We also thank Gopinath Krishnamoorthy and Kristine Arnvig for the helpful advice and critical reading of the manuscript. We acknowledge Caitlin Taylor for technical assistance with the CellASC platform and members of the MMRU for their advice and helpful discussions. This work was supported by grants from the Howard Hughes Medical Institute (Senior International Research Scholar’s grant to V.M.), the South African Medical Research Council (to V.M.) and the National Research Foundation of South Africa (to V.M. and D.F.W.).

## Data Availability

Raw fastq files for whole genome sequencing data for *M. smegmatis* mc^2^155, Δ*cobK*, and Δ*cobK* Δ*metH* are available in the European Nucleotide Archive under the accession numbers ERS3716042, ERS3716043, and ERS3716041, respectively. Supplementary Movies S1–5 can be accessed at https://uct.figshare.com/s/65105b9914196c4b4654.

---

**Table S1:** Strains and plasmids used in this study.

| Strain or plasmid | Description | References |
| --- | --- | --- |
| ***Mycobacterium smegmatis*** |  |  |
| **mc<sup>2</sup>155** | High-frequency transformation mutant of *M. smegmatis* ATCC 607 | (57) |
| **$\Delta cobK$** | *cobK* knock-out in mc<sup>2</sup>155 | This study |
| **$\Delta cobK\Delta methH$** | *methH* knock-out in $\Delta cobK$ | This study |
| **$\Delta metE cobK::hyg$** | *metE* knock-out and insertion of a *hyg* fragment in *cobK* | Dr. Stephanie Dawes, unpublished |
| ***methH* cKD** | *M. smegmatis methH* conditional knockdown strain | This study |
| **Plasmids** |  |  |
| **p2NIL** | Suicide plasmid; Km<sup>R</sup> | (46) |
| **pGOAL17** | Counter-selection cassette plasmid; Amp<sup>R</sup> | (46) |
| **pGOAL19** | Counter-selection cassette plasmid; Amp<sup>R</sup> | (46) |
| **pIJ963** | Hyg-cassette plasmid | (47) |
| **p3875K** | *M. smegmatis* $\Delta cobK$ vector; Km<sup>R</sup> | This study |
| **p3875K19** | *M. smegmatis* $\Delta cobK$ vector containing *PacI* cassette from pGOAL19; Km<sup>R</sup>, Hyg<sup>R</sup>, Suc<sup>S</sup> | This study |
| **p4185K** | *M. smegmatis* $\Delta methH$ vector; Km<sup>R</sup> | This study |
| **p4185K19** | *M. smegmatis* $\Delta methH$ vector containing *PacI* cassette from pGOAL19; Km<sup>R</sup>, Hyg<sup>R</sup>, Suc<sup>S</sup> | This study |
| **p4185K17** | *M. smegmatis* $\Delta methH::hyg$ vector containing *PacI* cassette from pGOAL19; Km<sup>R</sup>, Hyg<sup>R</sup>, Suc<sup>S</sup> | This study |
| **PLJR962** | CRISPRi backbone for *M. smegmatis* | (25) |
| **PLJR962_methH1** | *M. smegmatis* *methH* knock-down | This study |
|  | construct with sgRNA1 oligo |  |
| **PLJR962_methH2** | *M. smegmatis* *methH* knock-down<br>construct with sgRNA2 oligo | This study |
| **PLJR962_methH3** | *M. smegmatis* *methH* knock-down<br>construct with sgRNA3 oligo | This study |
| **PLJR962_methH4** | *M. smegmatis* *methH* knock-down<br>construct with sgRNA4 oligo | This study |
| **PLJR962_methH5** | *M. smegmatis* *methH* knock-down<br>construct with sgRNA5 oligo | This study |
| **PLJR962_methH6** | *M. smegmatis* *methH* knock-down<br>construct with sgRNA6 oligo | This study |
| **PLJR962_methH7** | *M. smegmatis* *methH* knock-down<br>construct with sgRNA7 oligo | This study |
| **PLJR962_methH8** | *M. smegmatis* *methH* knock-down<br>construct with sgRNA8 oligo | This study |
| **PLJR962_methH9** | *M. smegmatis* *methH* knock-down<br>construct with sgRNA9 oligo | This study |
| PLJR962_methH10 | *M. smegmatis* *methH* knock-down construct with sgRNA10 oligo | This study |
| PLJR962_methH11 | *M. smegmatis* *methH* knock-down construct with sgRNA11 oligo | This study |
| PLJR962_methH12 | *M. smegmatis* *methH* knock-down construct with sgRNA12 oligo | This study |
| PLJR962_methH15 | *M. smegmatis* *methH* knock-down construct with sgRNA15 oligo | This study |
| PLJR962_mmpL3 | *M. smegmatis* *mmpL3* knock-down construct | This study |

**Table S2:**
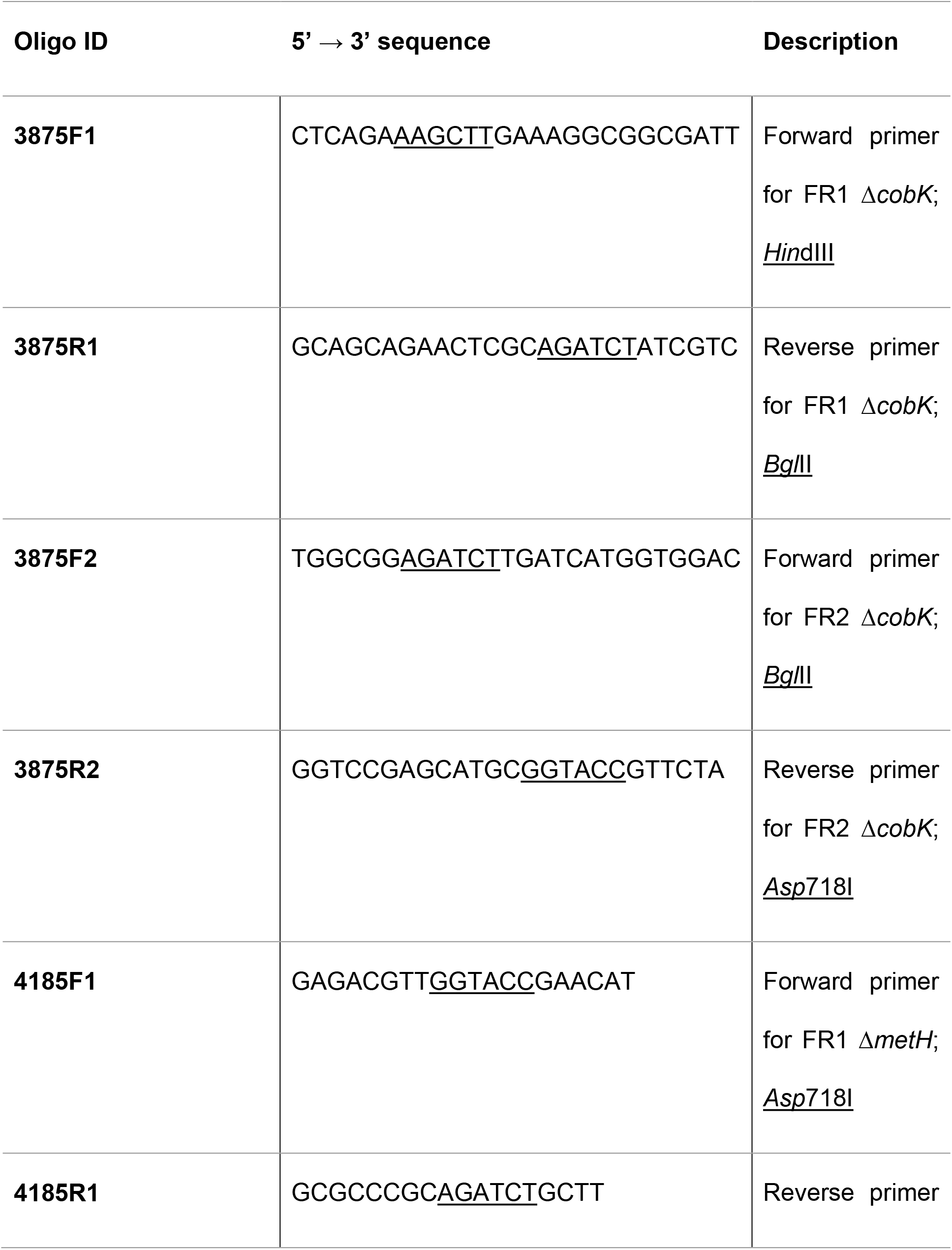
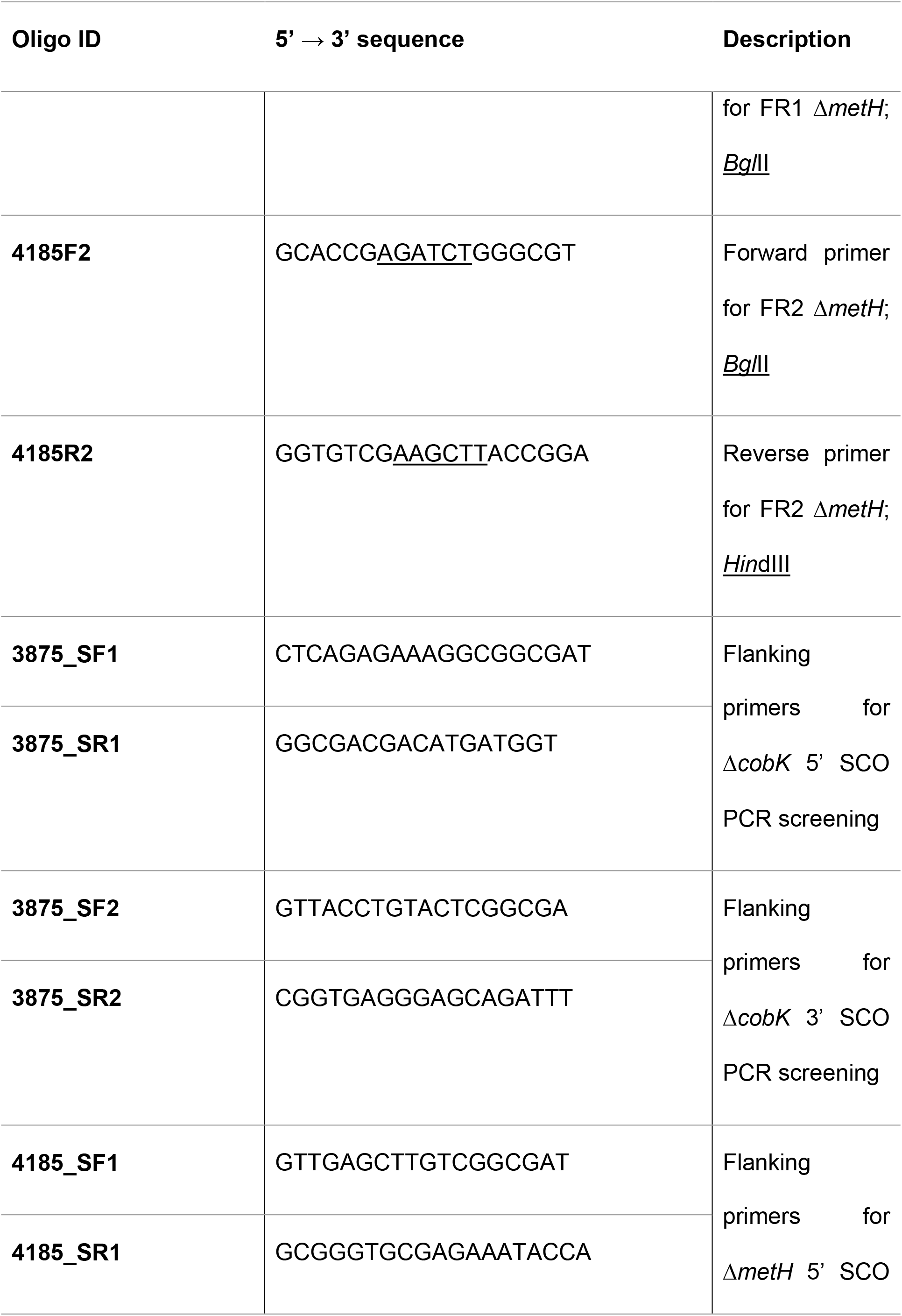
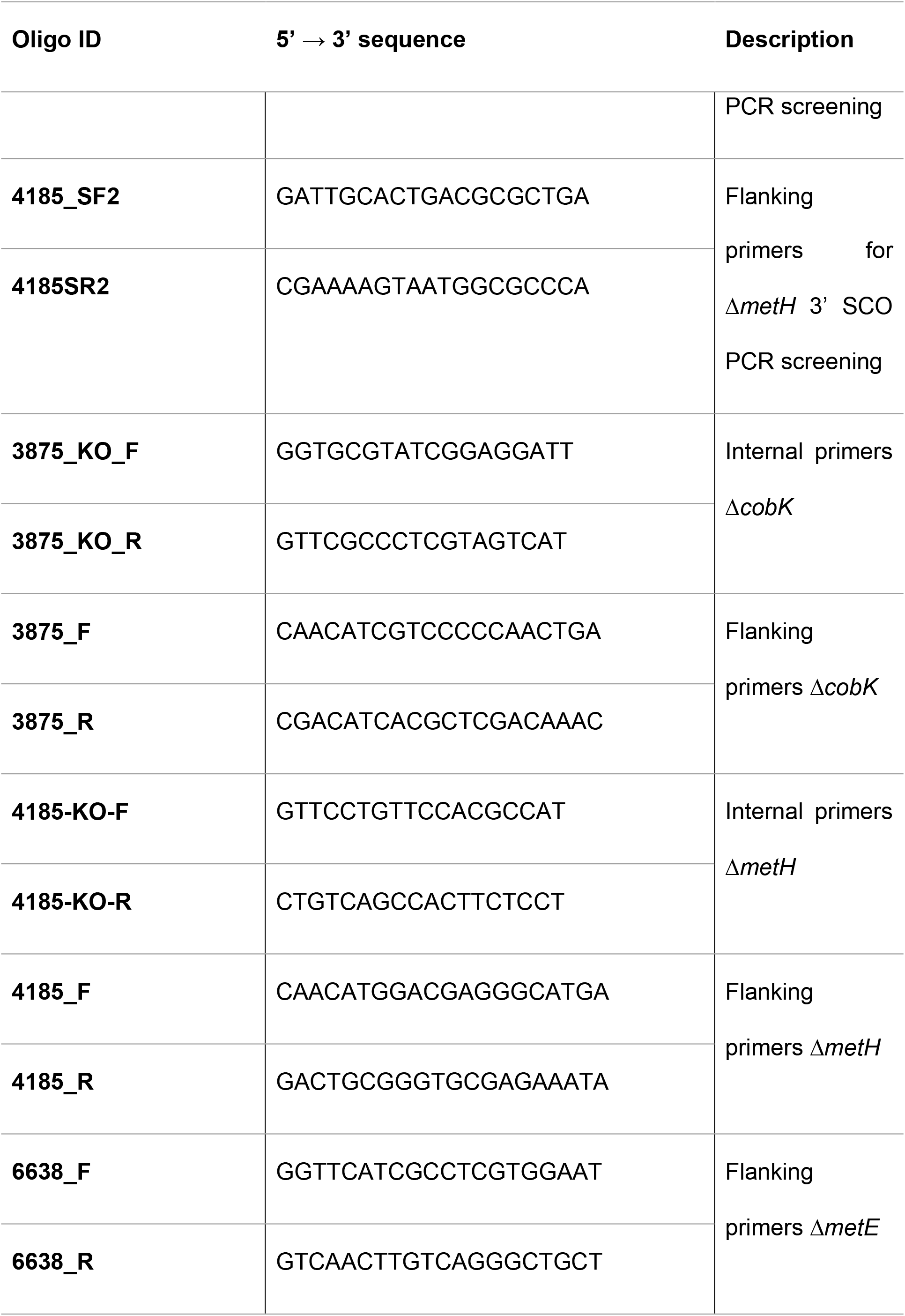
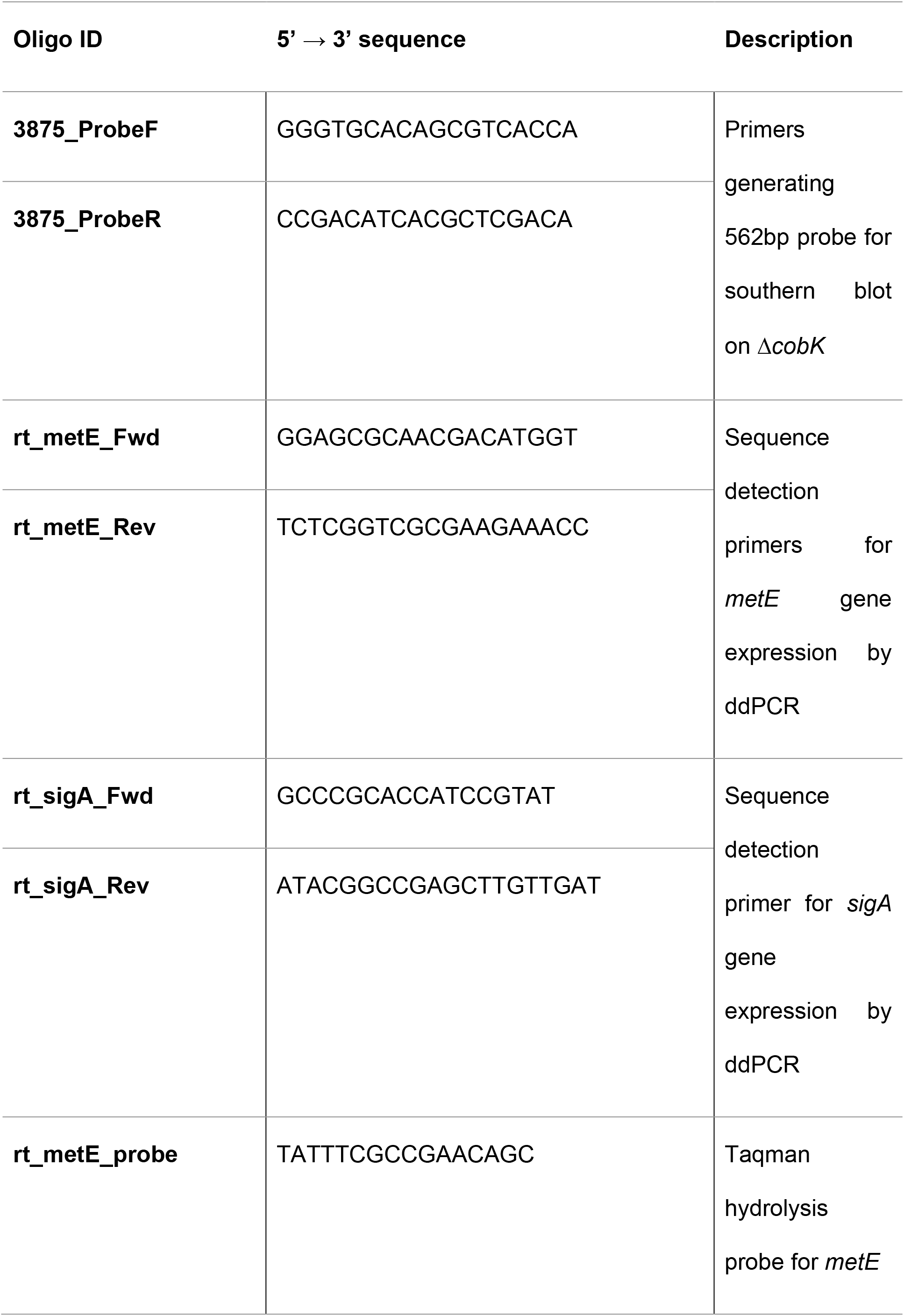
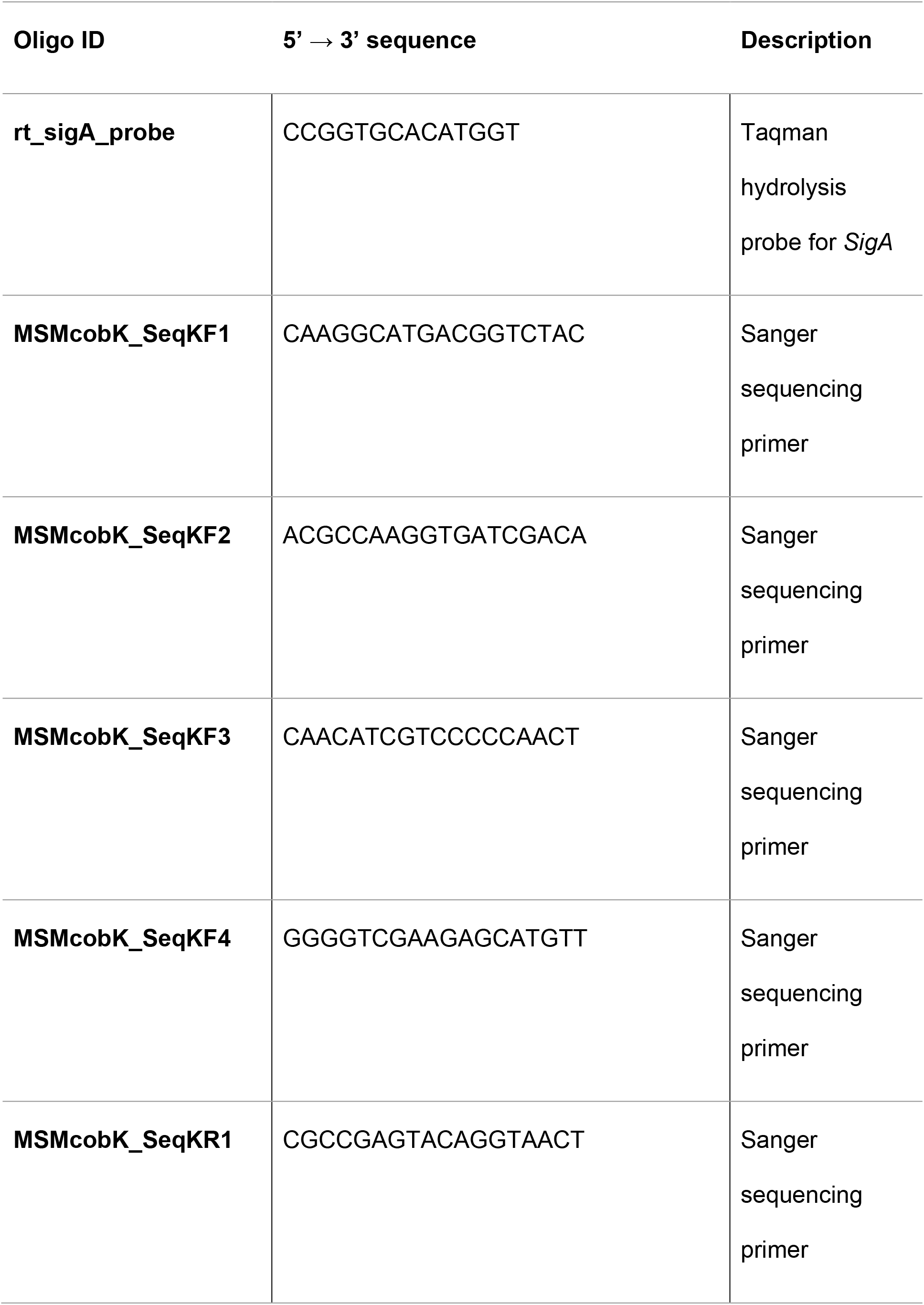
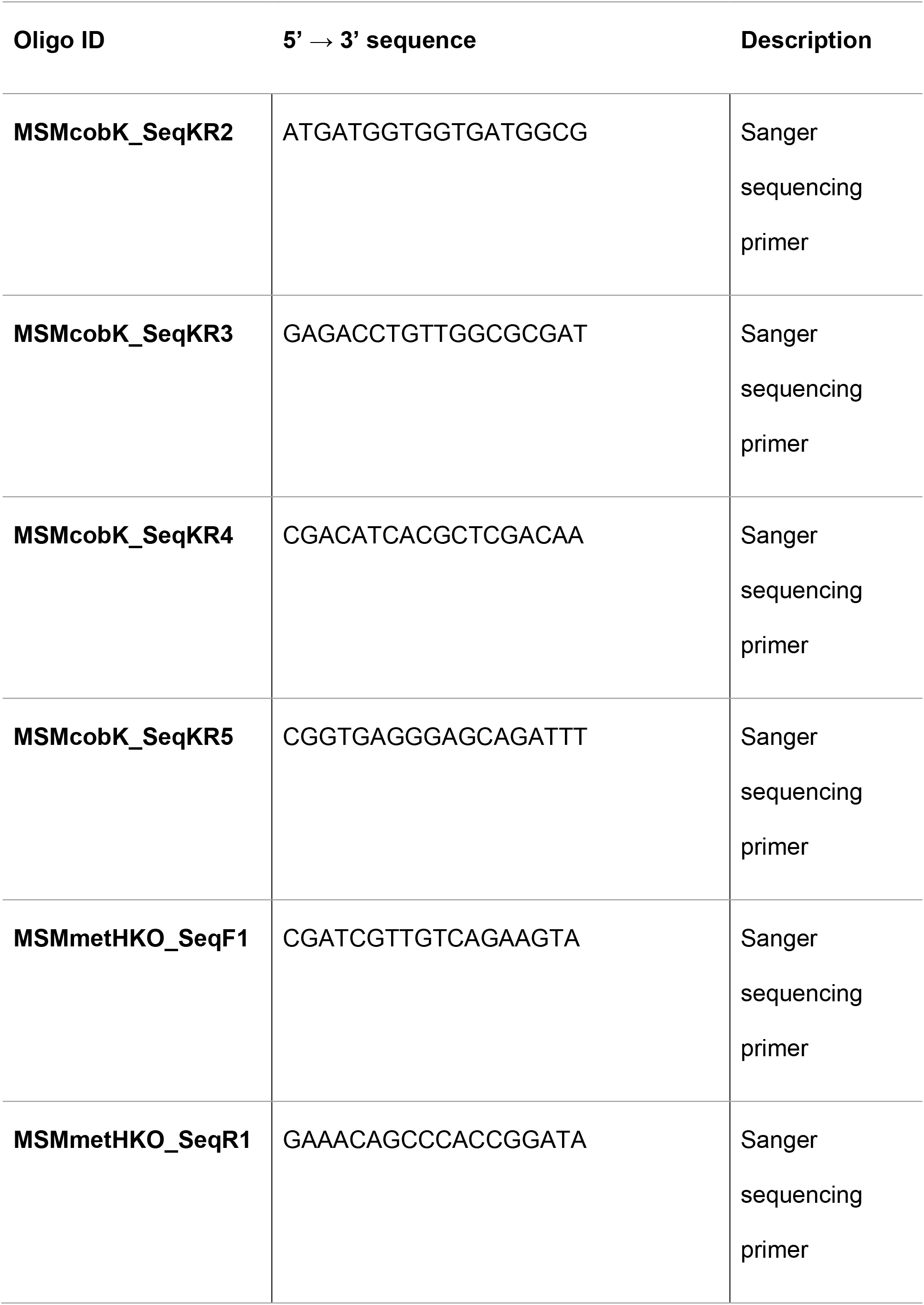
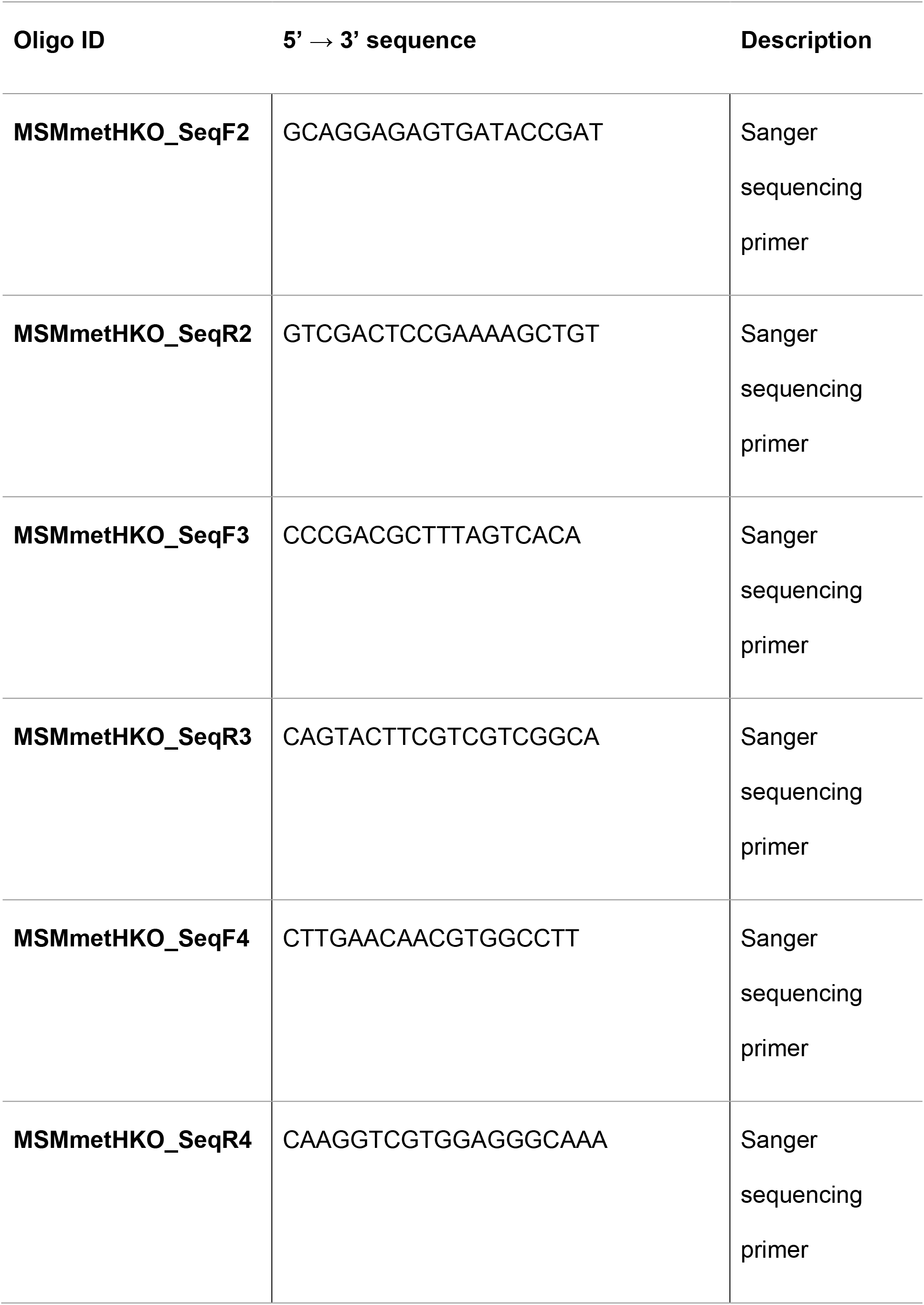
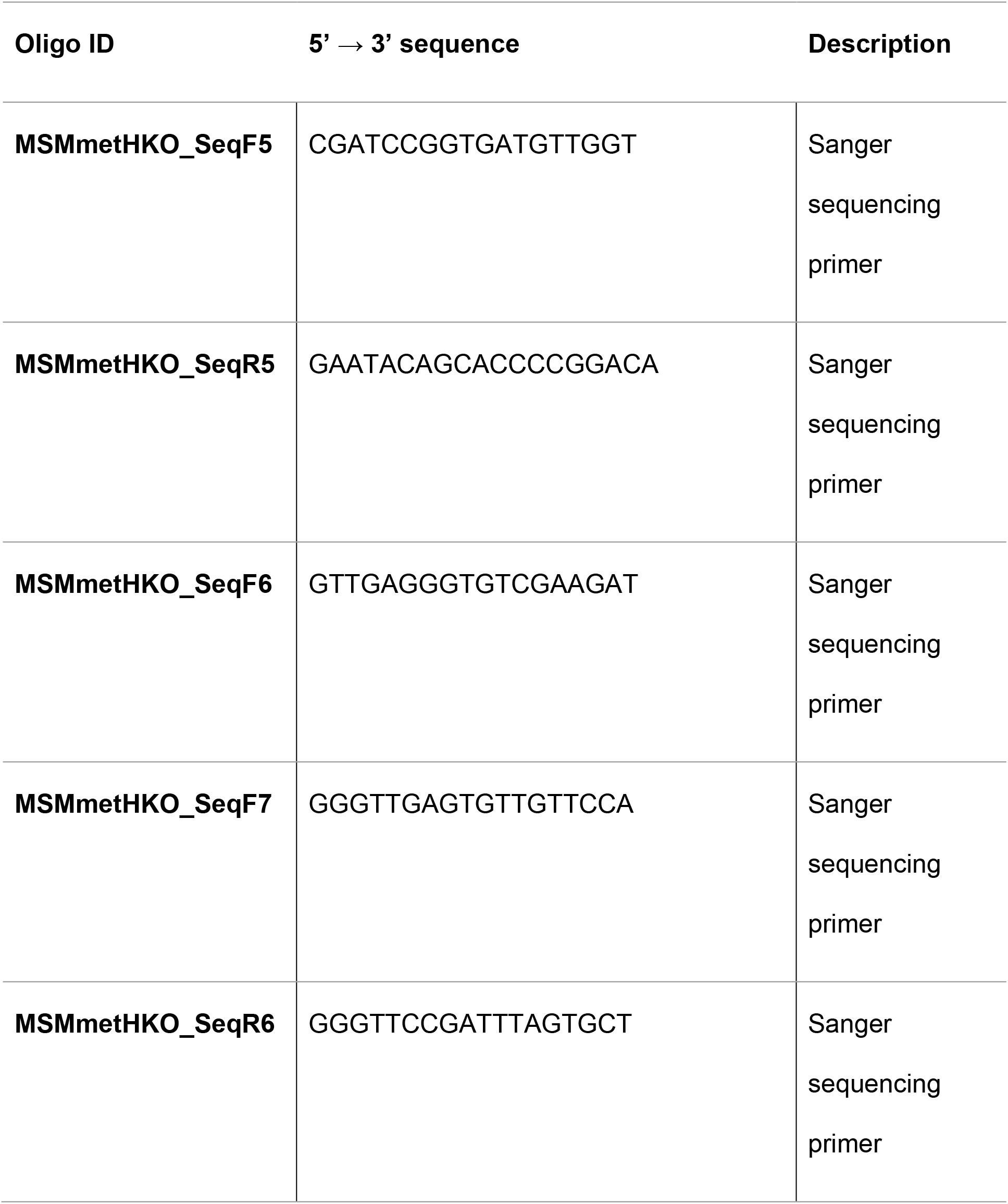
Oligos used for cloning, PCR screening, Sanger sequencing and gene expression analysis.

**Table S3:**
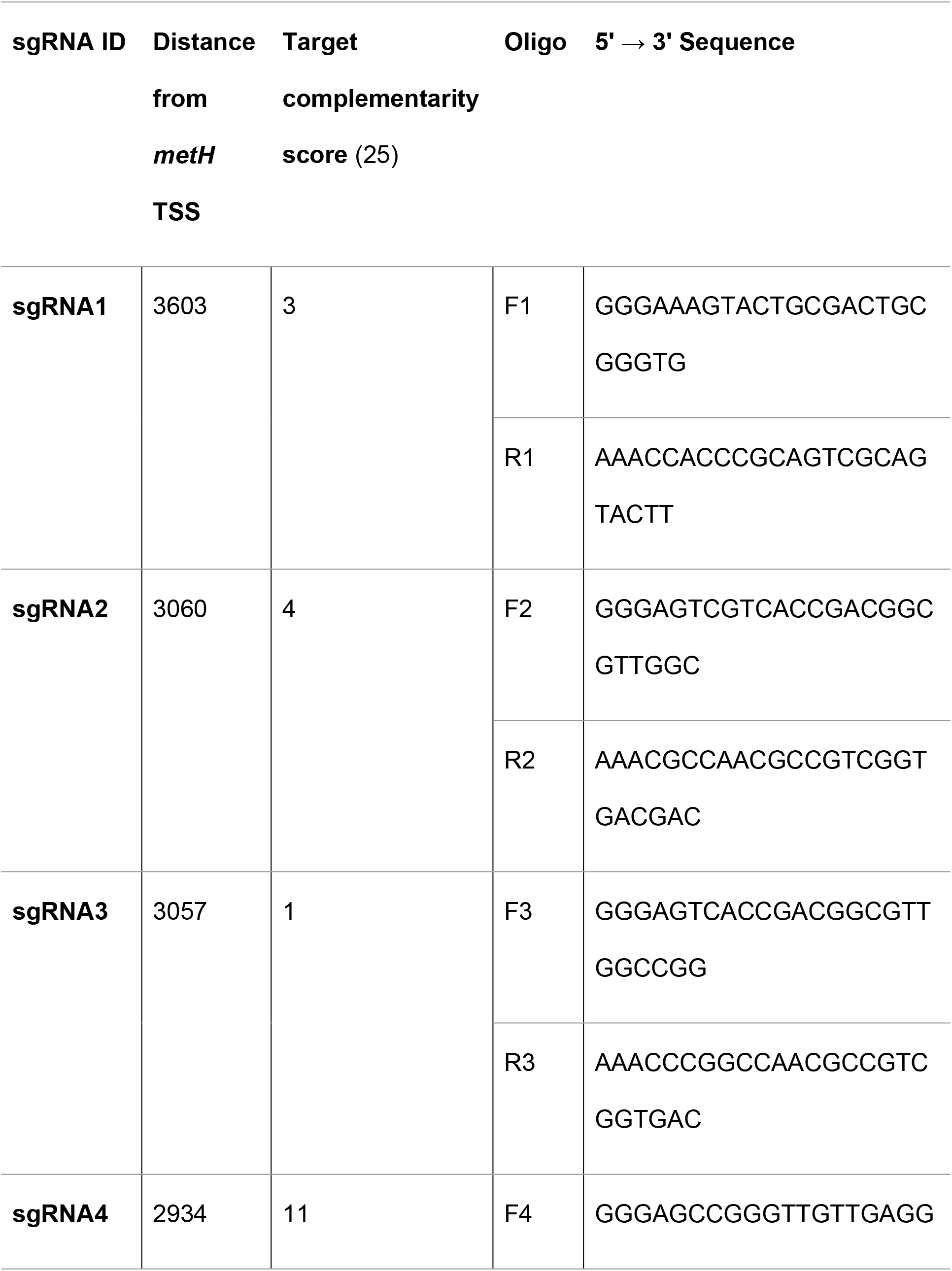
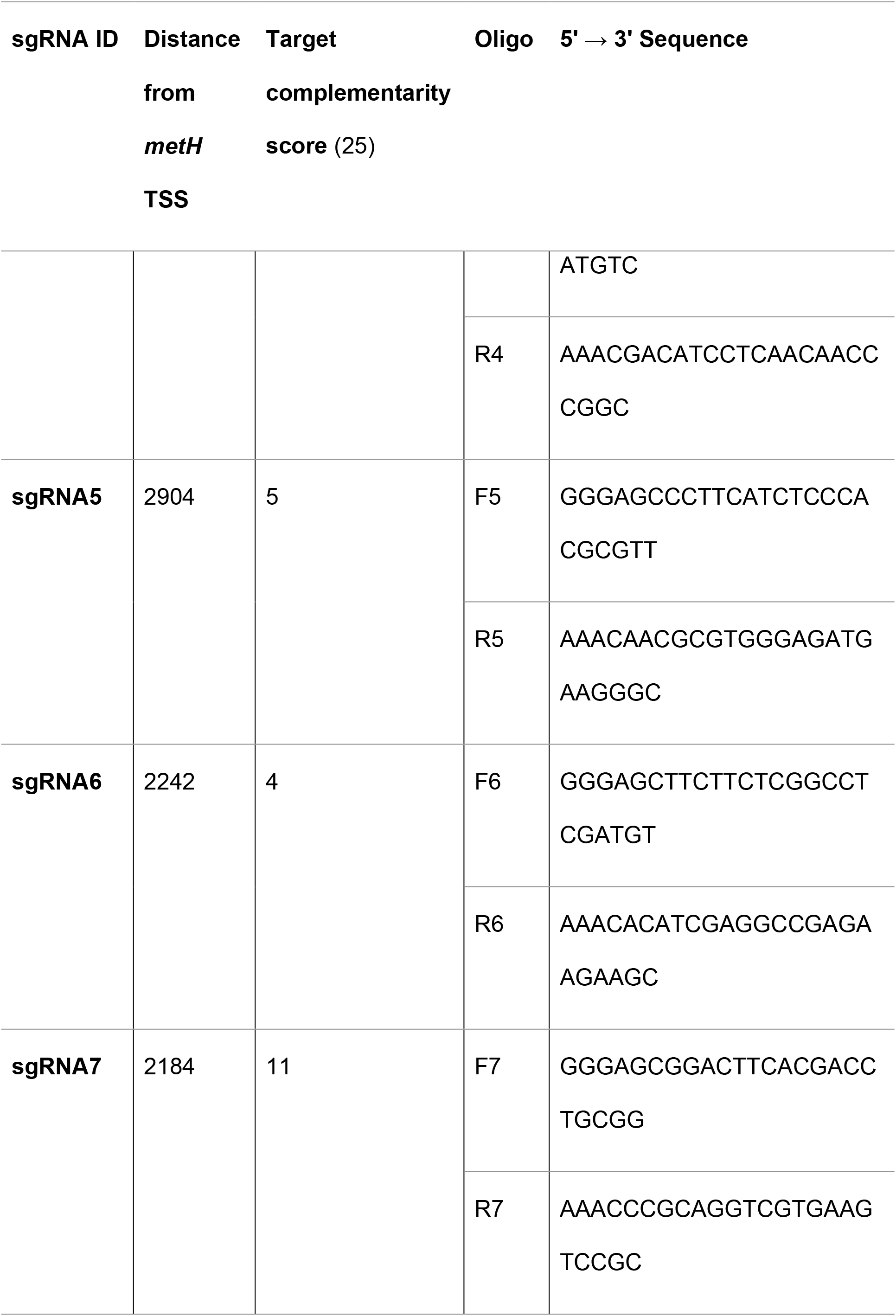
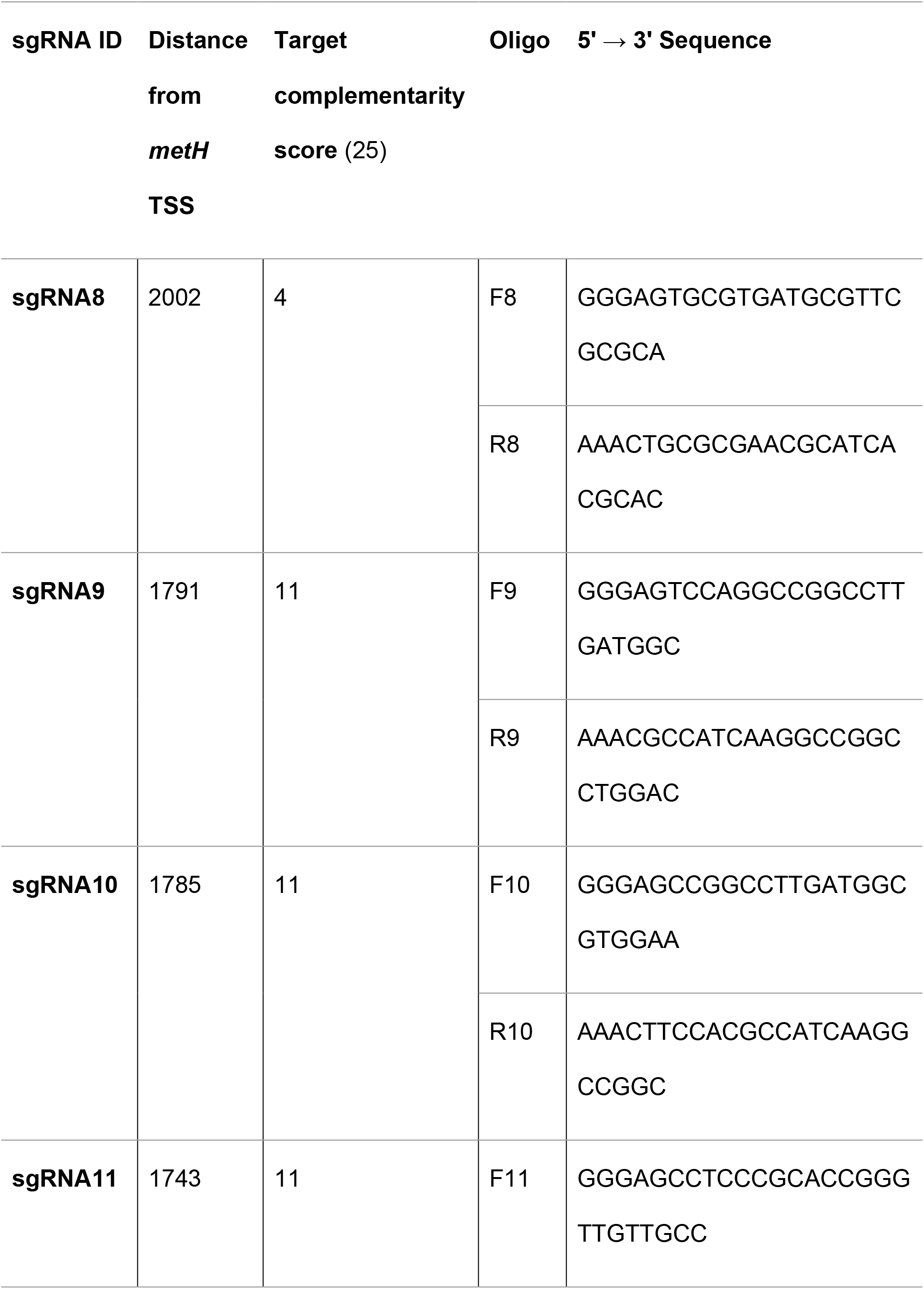
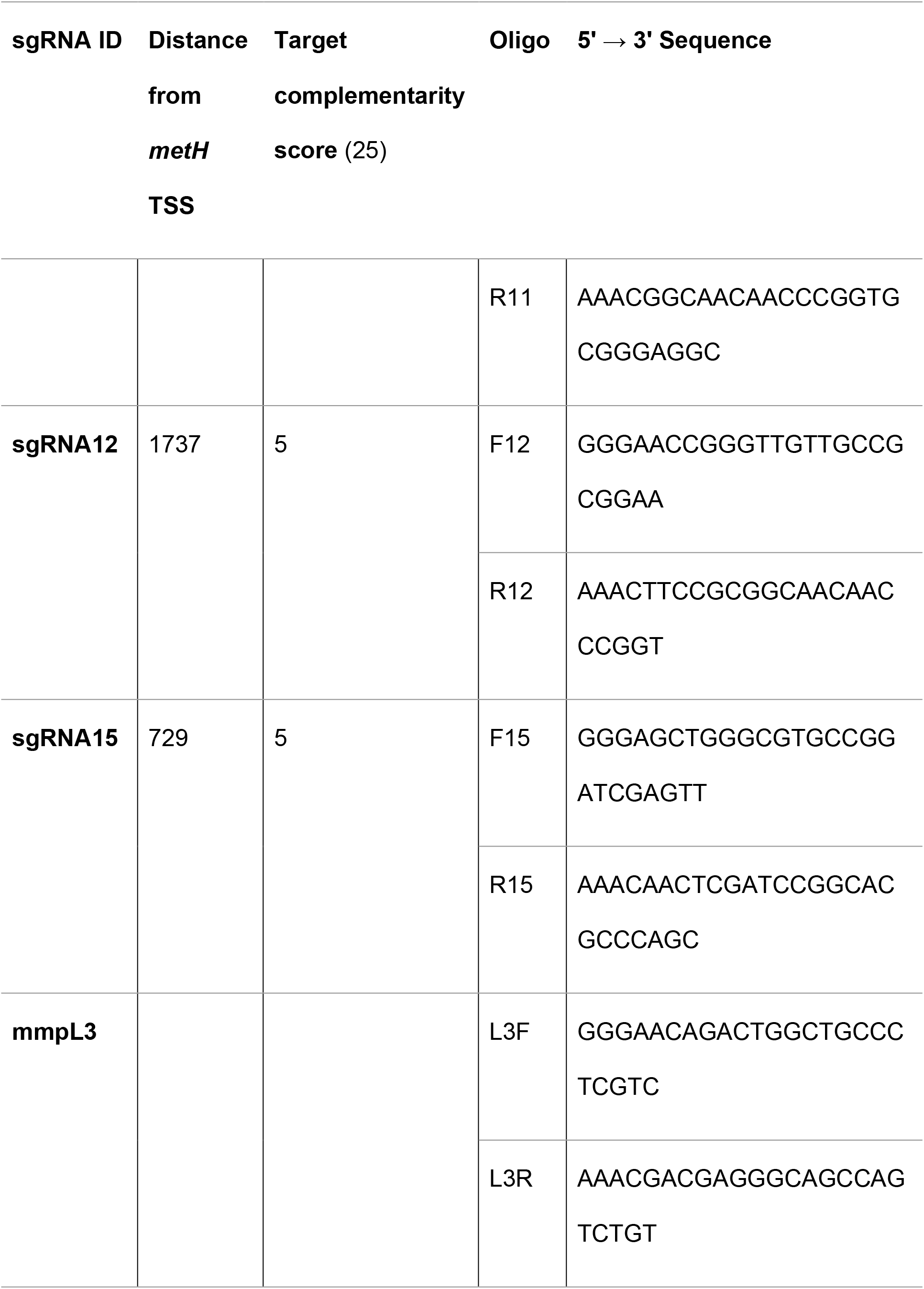
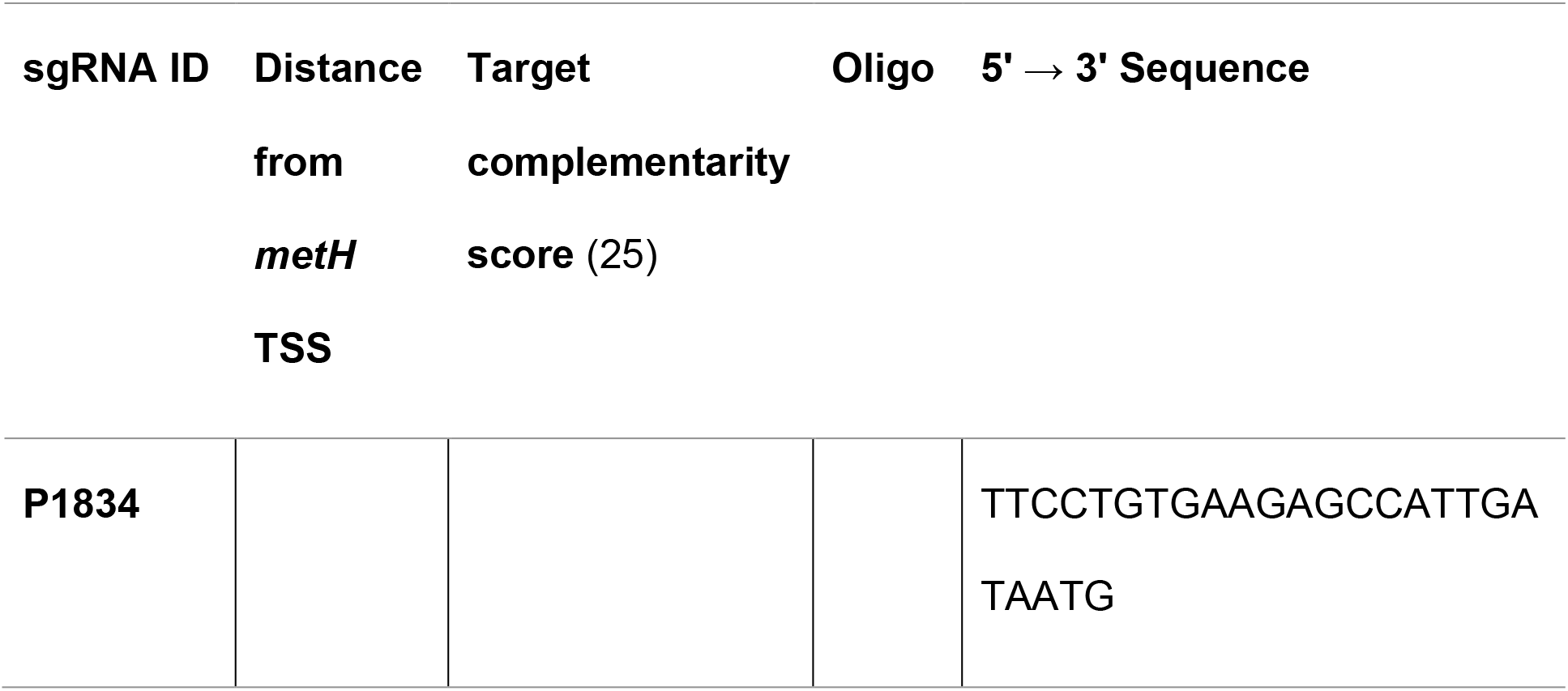
sgRNAs targeting *M. smegmatis metH* gene.

**Figure S1.**
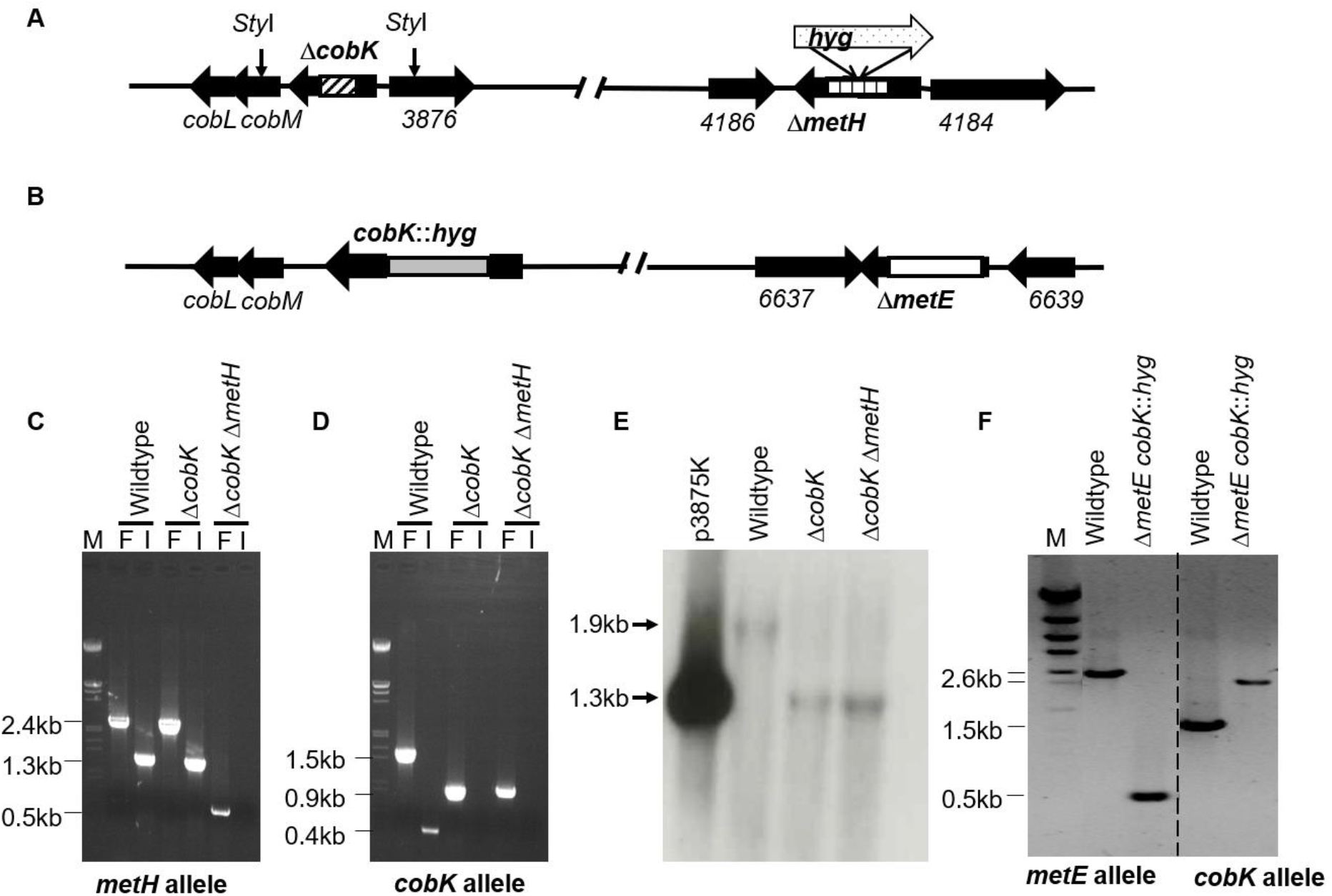
Construction and screening of *M. smegmatis* mutants. **A**. A schematic of the Δ*cobK* and Δ*metH* genotypes. The marked Δ*metH*::*hyg* construct was generated by inserting a *hyg* fragment (broad arrow with dotted pattern) was inserted into Δ*metH* construct using *Bgl*II sites. **B.** A schematic depicting the Δ*metE cobK*::*hyg* genotype showing the deleted portion of *metE* allele (white rectangle) and the insertion of a *hyg* fragment (grey rectangle) into the *cobK* allele. **C-D.** PCR screening of the putative Δ*cobK* and Δ*metH* strains using primers targeting flanking or internal (I) regions of the deleted portion of the genes. Amplicon sizes for flanking primers: wild type *metH* – 2.4kb; Δ*metH* – 0.5kb; wild type *cobK* – 1.5kb; Δ*cobK* – 0.9kb. Amplicon sizes for internal primers: wild type *metH* – 1.3kb; wild type *cobK* – 0.4kb. M – DNA molecular weight marker. **E**. Confirmation of Δ*cobK* by Southern blotting. A PCR-generated probe for *cobK* was used to detect a fragment between two naturally occurring *Sty*I restriction sites (down-facing arrows in **A**). Expected fragments: wild type *cobK* – 1.9kb; Δ*cobK* – 1.3kb. **F.** PCR genotyping of Δ*metE cobK*::*hyg* strain using primers flanking *cobK* and *metE*. Dashed line marks the border of two separate gels.

**Figure S2.**
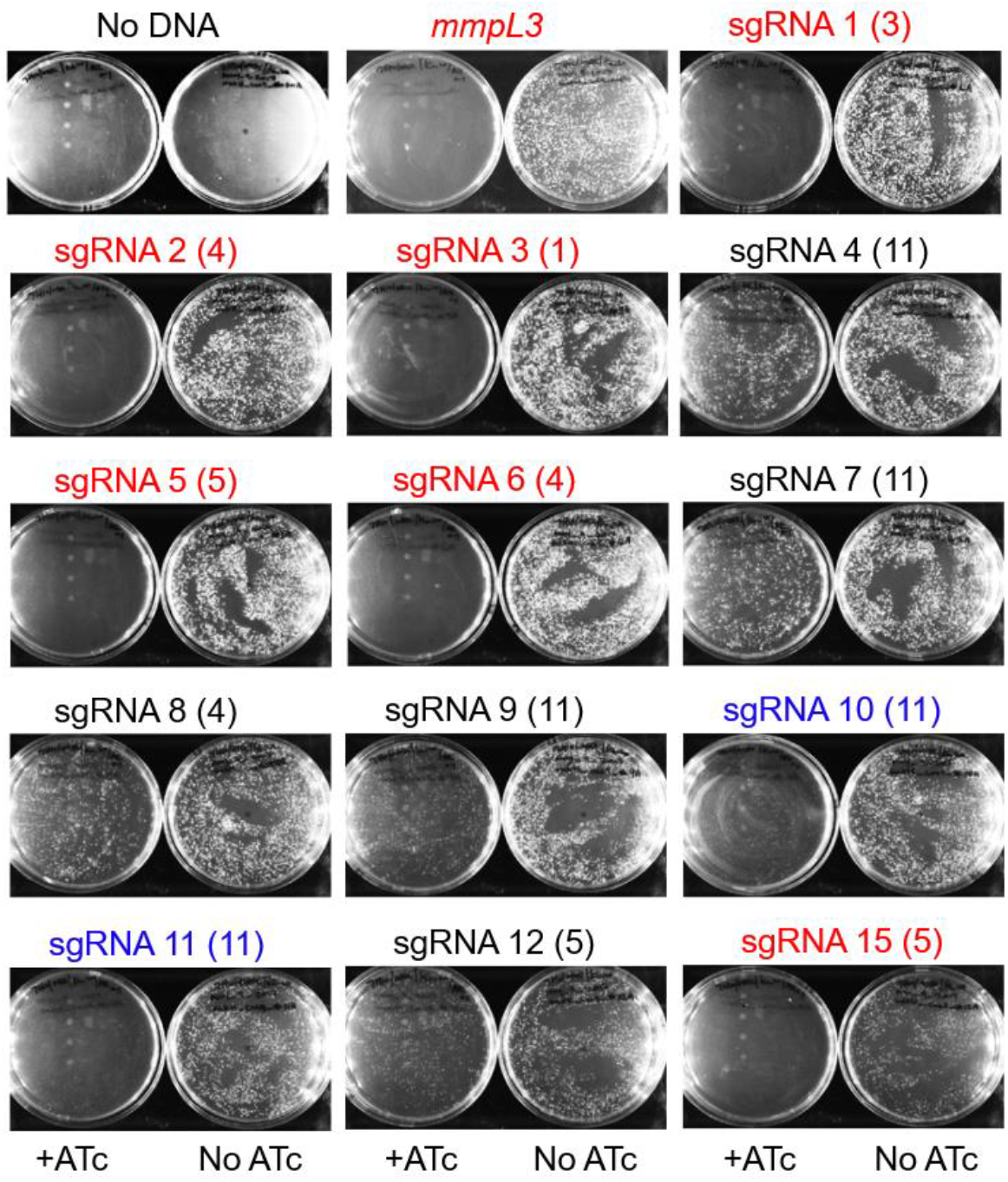
Conditional knockdown of *metH* using CRISPRi. The knock-down of *mmpL3* completely inhibited growth, whereas the 13 *metH* cKD constructs suppressed growth to varying degrees. The sgRNAs that produced complete inhibition of growth upon the ATc induction are highlighted in red, those exhibiting partial inhibition in blue, and those with no inhibition in black. The target complementarity scores associated with each sgRNA are indicated in parentheses.

## SUPPLEMENTARY MOVIES LEGEND

Live-cell imaging of *M. smegmatis* wild type and *metH* cKD strains using time-lapse phase-contrast microscopy. For single cell analysis using microfluidics, a suspension of 2 ×10^6^ bacterial cells/mL at exponential growth phase was preincubated with or without 100ng/mL of ATc for 6 h at 37°C prior to loading on the microfluidics platform. The experiment was run for 43 h and images were captured every 15 min.

